# Endogenous FGFs drive ERK-dependent cell fate patterning in 2D human gastruloids

**DOI:** 10.1101/2024.07.08.602611

**Authors:** Kyoung Jo, Zong-Yuan Liu, Gauri Patel, Zhiyuan Yu, LiAng Yao, Seth Teague, Craig Johnson, Jason Spence, Idse Heemskerk

**Affiliations:** Department of Cell and Developmental Biology, University of Michigan Medical School, Ann Arbor, Michigan; Department of Computational Medicine and Bioinformatics, University of Michigan Medical School, Ann Arbor, Michigan; Center for Cell Plasticity and Organ Design, University of Michigan Medical School, Ann Arbor, Michigan; Department of Biomedical Engineering, University of Michigan, Ann Arbor, Michigan; Department of Internal Medicine, Gastroenterology, University of Michigan Medical School, Ann Arbor, Michigan; Department of Physics, University of Michigan, Ann Arbor, Michigan

## Abstract

The role of FGF is the least understood of the morphogens driving mammalian gastrulation. Here we investigated the function of FGF in a stem cell model for human gastrulation known as a 2D gastruloid. We found a ring of FGF-dependent ERK activity that closely follows the emergence of primitive streak (PS)-like cells but expands further inward. We showed that this ERK activity pattern is required for PS-like differentiation and that loss of PS-like cells upon FGF receptor inhibition can be rescued by directly activating ERK. We further demonstrated that the ERK-ring depends on localized activation of basolaterally positioned FGF receptors (FGFR) by endogenous FGF gradients. We confirmed and extended previous studies in analyzing expression of FGF pathway components, showing FGFR1 is the main receptor, FGF2 is highly expressed across several cell types, and FGF4/17 are the main FGF ligands expressed in the PS-like cells, similar to the human and monkey embryo but different from the mouse. We found that knockdown of FGF4 greatly reduced PS-like differentiation while FGF17 knockdown primarily affected subsequent mesoderm differentiation. FGF8 expression was spatially displaced from PS-markers and FGF4 expression and peaked earlier, while knockdown led to an expansion in PS-like cells, suggesting FGF8 may counteract FGF4 to limit PS-like differentiation. Thus, we have identified a previously unknown role for FGF-dependent ERK signaling in 2D gastruloids and possibly the human embryo, driven by a mechanism where FGF4 and FGF17 signal through basally localized FGFR1 to induce PS-like cells and their derivatives, potentially restricted by FGF8.

## Introduction

During gastrulation the three germ layers are established, and the body plan is laid out. In amniotes this involves the formation of the primitive streak on the posterior side of the embryo from which cells ingress beneath the epiblast and migrate to form the mesodermal and endodermal layers. An important role in gastrulation is played by evolutionarily conserved morphogens from the BMP, Wnt, Nodal, and FGF families^1^. The function of the first three signaling pathways is comparatively well understood: in both mouse and human a transcriptional hierarchy between BMP4, Wnt3, and Nodal plays a central role in inducing dynamic signaling gradients along the anterior posterior axis that control cell fate patterning^1–3^. For BMP and Nodal, basal receptor localization in the epiblast is crucial to form signaling activity gradients^3–7^. Wnt and Nodal signaling are directly required for mesoderm and endoderm differentiation, while the balance between BMP and Nodal signaling is important for patterning the germ layers along the body axes^8^.

In contrast, the role of FGF in mammalian and especially human gastrulation has been more elusive. Secreted FGFs signal through FGF receptor tyrosine kinases (FGFRs) in a heparan sulfate proteoglycan (HSPG)-dependent manner to activate several signaling pathways including MAPK/ERK, PI3K, and PLCγ^9^. There is evidence for a conserved requirement for FGF/ERK signaling in mesoderm formation across vertebrate model organisms^10–13^. However, it is unknown if there are FGF- and phosphorylated ERK gradients across the mammalian primitive streak^14,15^, whether there are functionally relevant ERK signaling dynamics like in other contexts, or whether receptor localization is important for establishing signaling gradients^16–19^. It is also unclear to what extent the requirement in mesoderm induction is direct, or indirect by modulating other signals^14,15,20^. In addition, several FGFs are co-expressed and the roles of specific FGFs are uncertain. Moreover, there may be significant interspecies differences in this respect. Notably, FGF8 is the only FGF which has been shown to be required for gastrulation in the mouse^21^, and its requirement in gastrulation appears to be conserved across vertebrate model organisms^22–24^, yet FGF8 is not highly expressed in primate embryos^25,26^.

Mutants for FGF8, FGFR1, HSPG synthesis, and ERK2, as well as pharmacological inhibition of MEK and FGFR give rise to similar phenotypes in mouse gastrulation, suggesting that FGFR1 and FGF8 are the only FGF ligand-receptor pair required for gastrulation and that these act through the MAPK/ERK pathway^12,20,21,27–31^. FGF4 expression during gastrulation is also lost in FGF8 mutants, leaving the possibility of a role for FGF4, but this remains unknown as FGF4 is also required in the blastocyst and mutants therefore do not reach gastrulation^32^. In all mutants and for pharmacological inhibition of FGF/ERK signaling, expression of the primitive streak marker brachyury (TBXT, also BRA, or T in mouse) is lost or severely diminished in the posterior streak and mesoderm fails to migrate away, piling up in this region. In addition, TBX6 expression is lost completely, and no lateral or paraxial mesoderm are formed. In contrast, TBXT expression in the anterior streak and extraembryonic mesoderm is not dependent on FGF/ERK signaling; extraembryonic mesoderm development is normal, and the axial mesoderm is expanded^20,21,27,28^.

In vitro models offer an opportunity to disentangle the complexity of FGF signaling in human development. Here, we explore the role of FGF/ERK signaling in micropatterned human pluripotent stem cells treated with BMP4, also known as 2D human gastruloids. These give rise to concentric rings of different cell types associated with gastrulation after 2 days of differentiation. Pluripotent cells in the center are surrounded by a primitive streak-like ring, which itself is surrounded by primordial germ cells and amnion-like cells on the colony edge^33–35^. Due to its simplicity and reproducibility, this system has proven powerful in deciphering the mechanisms underlying aspects of human gastrulation. It was used to show that BMP-Wnt-Nodal hierarchy that controls primitive streak formation during mouse gastrulation is preserved in human while providing a range of new insights into how these signals function^2,3^. For example, the crucial role for BMP and Nodal receptor localization in patterning was discovered in 2D gastruloids and later confirmed in the mouse embryo. This system also revealed that BMP, Wnt, and Nodal signaling patterns are highly dynamic^3–7^. It therefore also provides a promising approach to determine how FGF may function in human gastrulation.

In the 2D human gastruloid model, we show that a ring of phosphorylated ERK forms in an FGF-dependent manner and expands largely in lockstep with the formation of primitive streak-like cells but extends further inward. We demonstrate that this pattern of ERK activity is due to localized FGF receptor (FGFR) activation that requires basal FGFR localization. We then show that exogenous FGF2 in the media is required for pattern formation to occur but can be removed well before the phosphorylated (active) ERK (pERK) and PS-like rings appear, suggesting the pERK ring instead depends on endogenous FGF gradients. Using single cell RNA-sequencing, we determine the expression of FGF pathway components including FGFR1 as the main receptor and FGF2, FGF4, and FGF17 as the most highly expressed ligands. Our analysis corroborates previous studies describing expression of FGF2 and FGF17 in human embryos and gastruloids but for the first time recognizes a possible role for FGF4 and includes a broad analysis of genes involved in FGF signaling modulation^25,36^. We find that the FGF4 and FGF17 expression are restricted to the PS-like ring and that knockdown of FGF4 significantly reduces PS-like differentiation, while FGF17 knockdown primarily affects downstream mesoderm differentiation. Given its importance in the mouse, we also analyze FGF8 and find its expression is spatially and temporally displaced from FGF4 and PS-markers, while FGF8 knockdown leads to an increase in PS-like differentiation, suggesting it may counteract FGF4. In summary, we have greatly advanced our understanding of how FGFs function in a model for human gastrulation.

## Results

### FGF/ERK signaling is required for primitive streak-like differentiation

We previously observed that pharmacological inhibition of MEK (the kinase upstream of ERK) or FGF receptors leads to a loss of differentiation to primitive streak-like and primordial germ cell-like cells, suggesting a crucial role for FGF/ERK signaling^34^. Moreover, a ring of active ERK (pERK) has been observed in 2D gastruloids but its connection to FGF signaling or differentiation remains unclear^33^. Therefore, we wanted to investigate the role of FGF/ERK signaling further.

To determine the dynamics of pERK and relate it to PS-like differentiation, we first performed time series immunofluorescence co-staining for pERK and the PS marker TBXT after BMP4 treatment in 700um diameter colonies. This revealed the pERK ring first emerges around 24h, around the same time as TBXT expression, and gradually gets brighter and wider (Fig. 1a). By 42h the outer edge of the ERK ring coincided with TBXT expression, while the inner edge was less sharp and extended further towards the center than TBXT, reminiscent of the domains of WNT and NODAL signaling that extend further in than the PS-like region^3,6^ (Fig. 1a). Cell fate patterning in 2D gastruloids has a fixed length scale from the edge, with the central part absent in smaller colonies due to the fixed length scale of the BMP signaling gradient from the edge. Consistently, we found that ERK signaling also has a fixed pattern from the edge regardless of colony size, so that in smaller colonies the ERK ring becomes a domain of high ERK signaling extending through the colony center (Supp. Fig. 1a). MEK or FGFR inhibition for 30 minutes after 41.5h eliminated ERK activity, suggesting ERK activity depends entirely on FGF and that continuous FGF signaling is necessary to maintain the ERK signaling pattern (Fig. 1bc). Inhibition for the full duration of differentiation eliminated TBXT expression as well as other PS markers (Fig. 1bc, Supp. Fig. 1b). However, we were able to rescue TBXT expression by directly and uniformly activating ERK (Fig. 1d-f, Supp. Fig. 1cd) in cells stably expressing the doxycycline-inducible membrane-targeted catalytic domain of SOS (dox-SOS^cat^)^37^. Despite uniform induced ERK activity, TBXT expression remained excluded from the center, showing that ERK activation alone is not sufficient for TBXT induction. However, TBXT levels were much higher in the rescue (Supp. Fig. 1cd), possibly reflecting the much higher level of ERK activity induced by dox-SOS^cat^. MEK and FGFR inhibition also led to a reduction in cell number which staining for BrdU and cleaved Cas3 suggested was primarily due to reduced proliferation (Supp. Fig. 1ef). In contrast to TBXT expression, reduced cell number caused by FGFR inhibition was only partially rescued by dox-SOS^cat^, indicating additional downstream pathways regulate cell number (Fig. 1e). Altogether, these data show the formation of a PS-like ring is dependent on FGF signaling through ERK.

**Figure 1.**
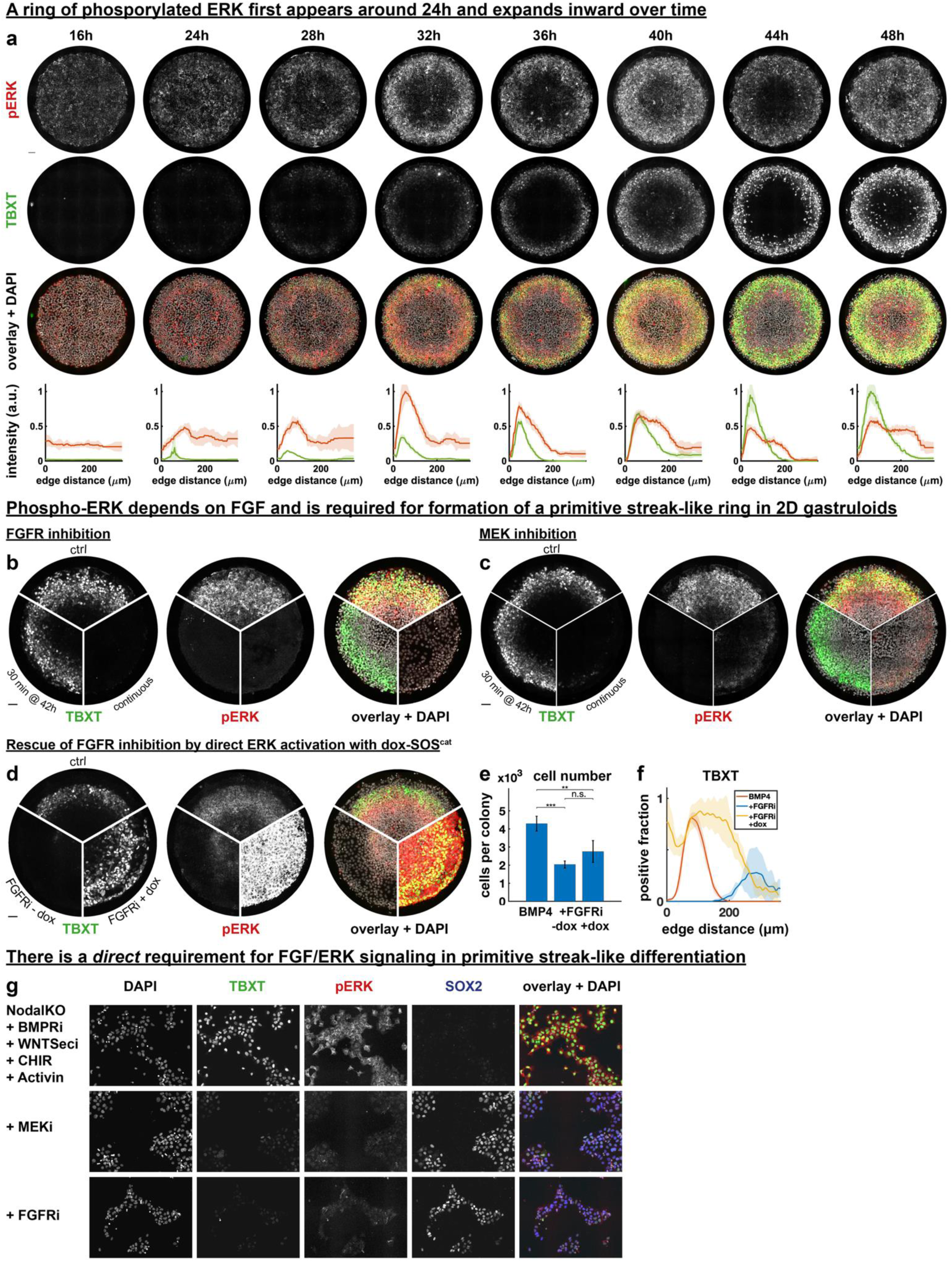
**a)** Immunofluorescence stainings for pERK and TBXT and their radial intensity profiles in 2D gastruloids at different times. Error bars in graphs are standard deviations over N=4 colonies. **b,c)** Effect FGF receptor inhibition (FGFRi, b) or MEK inhibition (MEKi, c) on pERK and TBXT in colonies fixed at 42h after BMP treatment. Inhibition for either 30 minutes at 41.5h (rounded to 42 in figure) or throughout differentiation. **d)** BMP only control and FGFRi inhibition in cells expressing doxycycline-inducible SOS^cat^ with and without addition of doxycycline. **e-f)** Cell numbers (e) and radial profiles of TBXT positive cells (f) for conditions in (d). Error bars represent standard deviation. Statistical significance was assessed using one-way ANOVA followed by Tukey’s HSD test for pairwise comparisons. **g)** PS marker TBXT, pERK and pluripotency marker SOX2 in Nodal knockout cells with inhibitors of Wnt secretion (WntSeci) and BMP receptors (BMPRi) with or without MEKi or FGFRi. All scale bars 50um.

This raised several questions. First, is FGF/ERK signaling required directly for PS-like differentiation, or does it act indirectly? These possibilities are not mutually exclusive. For example, FGF/ERK could be required directly but also act indirectly by controlling Wnt or Nodal expression, as both Wnt and Nodal signaling are required for PS-like differentiation. Second, which are the FGFs activating the ERK pathway? The pathway could be activated by exogenous FGF2 present in the pluripotency medium mTeSR1 or by endogenously expressed ligands. Third, how is ERK activity restricted to a ring?

To determine if there is a direct requirement of FGF/ERK signaling and whether this depends on exogenous FGF, we performed differentiation to PS-like cells in standard culture by exogenous activation of Wnt and Nodal signaling, bypassing the need to form endogenous Wnt and Nodal gradient as in 2D gastruloids^38^. We used a minimal base medium (E6) without exogenous FGF or other growth factors. To rule out any additional requirement for endogenous Nodal and Wnt which may be FGF/ERK-dependent, we used Nodal knockout cells^3^ treated with the Wnt secretion inhibitor IWP2, thereby eliminating any endogenous activity of these pathways. We also inhibited BMP receptors using LDN193189 to rule out any indirect effect through endogenous BMP. Under these conditions, with exogenous Wnt and Nodal activation for 24h in the absence of exogenous FGF, cells efficiently differentiated to TBXT+ PS-like cells (Fig. 1g, Supp. Fig. 1g). These PS-like cells had high active ERK levels just as in the 2D gastruloids. Inhibition of either FGFR or MEK abrogated ERK activity and TBXT expression while FGFR inhibition also severely reduced proliferation. In both cases significant expression of the pluripotency and ectoderm marker SOX2 was maintained (Supp. Fig. 1g). Given that FGF/ERK is known to be involved in pluripotency maintenance, this suggests inhibition may not be complete and that much lower ERK activity is required for pluripotency maintenance than PS-like differentiation, consistent with lower ERK activity in the pluripotent center of gastruloids.

MEK and FGFR inhibition reduce proliferation and thereby cell density, raising the possibility that their lack of differentiation is an indirect effect that depends on cell density: a critical density may be required for differentiation due to a community effect. To rule this out we performed the experiment across a range of densities, which confirmed that PS differentiation in the control was efficient even at the lowest densities while FGFR and MEK inhibition effectively blocked differentiation when compared to controls with similar final cell densities (Supp. Fig. 1h). We also tested whether proliferation and differentiation require similar levels of ERK signaling by determining the MEKi dose response of gastruloids. This revealed that lower doses of MEKi effectively block differentiation without strongly affecting cell number, further supporting that the effects of ERK signaling on proliferation and differentiation are independent (Supp. Fig. 1i). Altogether, our results show that endogenous FGF expression is both necessary and sufficient for PS-like differentiation by exogenous Wnt and Nodal stimulation and demonstrate a direct requirement for ERK signaling in PS-like differentiation.

### Single cell RNA-sequence reveals differentially expressed FGF pathway components

The above results suggest endogenous FGF expression may contribute to elevated ERK signaling in the PS-like ring. However, FGF/ERK signaling may be modulated at many levels (Fig. 2a). A spatial pattern of ERK activation could reflect a concentration gradient of FGF. Alternatively, differential expression of receptors and HSPG-modifying enzymes could lead to spatially patterned receptor activation and downstream ERK activity even in the absence of an FGF gradient. Even if receptor activation is uniform, downstream ERK activity could be spatially patterned by downstream negative regulators of ERK activity such as Sprouty-family proteins (SPRY) and dual-specificity phosphatases (DUSPs)^40^.

**Figure 2.**
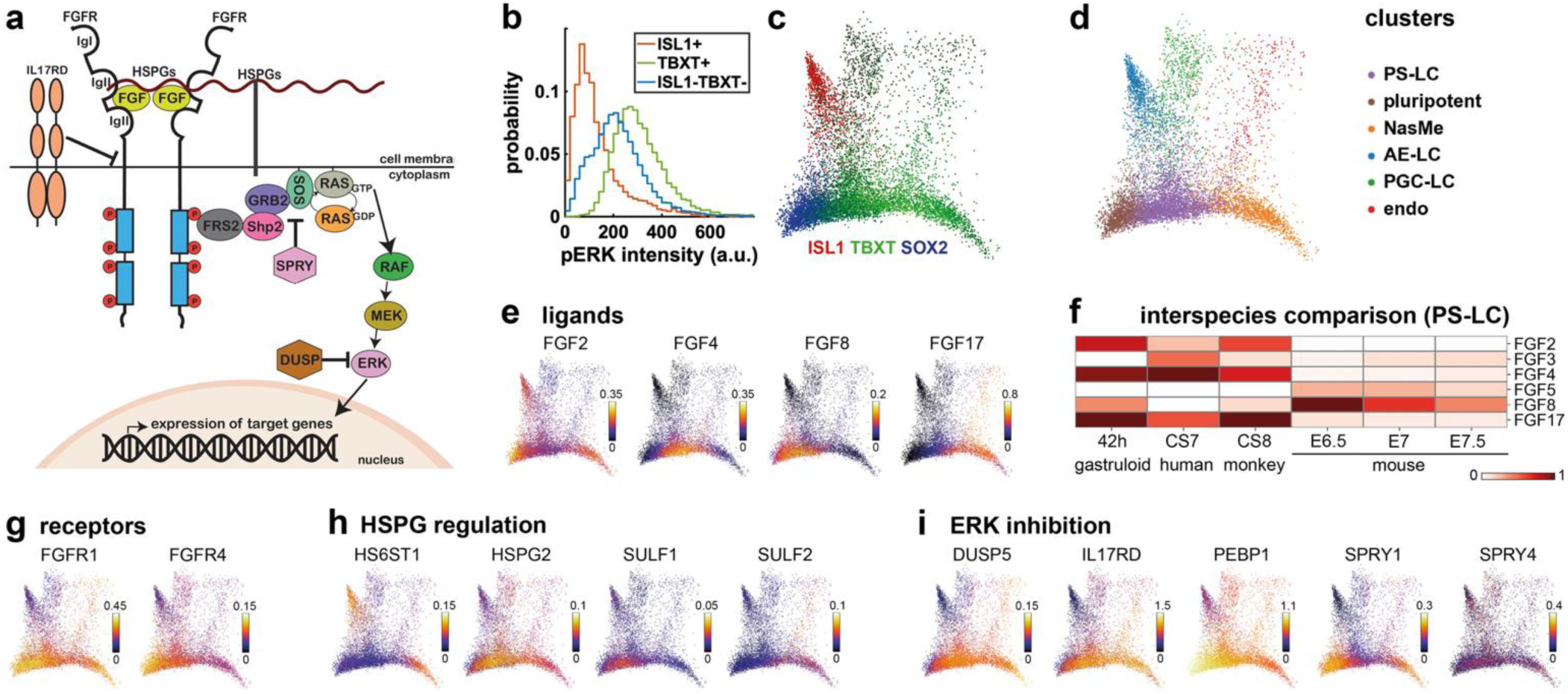
**a)** Schematic of the FGF/ERK signaling pathway. **b)** Distribution of pERK levels in TBXT+ cells versus ISL1+ and other cells based on data and thresholds in Supp. Fig. 2b. **c)** Expression of key markers in PHATE^39^ projection of scRNA-seq data for 42h colony. **d)** Cell type annotation. PS-LC: primitive streak-like cells, NasMe: nascent mesoderm, AE-LC: amniotic ectoderm-like cells: PGC-LC: primordial germ cell-like cells, endo: definitive endoderm. **e)** Expression of canonical FGF ligands (log transformed and smoothened). **f)** Comparison of FGF ligand expression in 2D human gastruloids to human, monkey, and mouse embryo, normalized within each type of sample (see methods). **g-i)** Expression of several genes involved in FGF signaling with strong differential expression between PS-like cells or nascent mesoderm and pluripotent cells, split by gene category.

To understand which genes modulating FGF/ERK signaling could be responsible for the elevated ERK signaling in TBXT positive cells and required for TBXT expression (Fig.1, Fig. 2b), we used single cell RNA-sequencing data from one previously published sample^34^ and one newly collected replicate (Supp. Fig. 2a) to determine expression of FGF pathway components in human 2D gastruloids, extending earlier work describing FGF expression^25,35,36,41^. TBXT-positive PS-like cells give rise to nascent mesoderm expressing TBX6 as well as definitive endoderm expressing SOX17 and FOXA2, both of which may still express TBXT at 42h^42^. To understand how *TBXT* expression is initiated in an FGF-dependent manner, we focused on differential expression in the PS-LCs relative to pluripotent cells, reasoning that FGF modulation specific to nascent mesoderm or endoderm is a consequence rather than a cause of PS-LC differentiation. We identified different cell types based on marker genes with the PS-like cells defined as *TBXT+MIXL1+POU5F1+TBX6-SOX17-FOXA2-*(Fig. 2cd, Supp. Table 4). We then analyzed expression of different categories of FGF/ERK regulators.

We found four FGF ligands with significant expression: *FGF2, 4, 8,* and *17* (Fig. 2e, Supp. Fig. 2b). We then compared the expression of these ligands in PS-like cells to reference data of CS7 human, CS8 monkey, and E6.5-E7.5 mouse gastrulation (Fig. 2f, Supp. Fig. 2cd). We also included *FGF3* and *FGF5*, which are expressed during mouse gastrulation^32,43,44^. High expression of *FGF4* and *FGF17* and low expression of FGF8 was consistent between human, monkey, and 2D gastruloid. *FGF2* expression was also high in the PS-like populations in 2D gastruloids and the CS8 monkey embryo, but much lower in the human embryo data. Furthermore, the monkey and human embryos expressed *FGF3*, which was minimally expressed in gastruloids. A similar comparison in nascent mesoderm showed much lower *FGF2/4* and much higher *FGF17* compared to PS across all primate samples (Supp. Fig. 2e). Expression in 2D gastruloids was consistent with published RNA-seq data for hPSC-derived PS-like cells, which expressed high *FGF2* and *FGF4*, some *FGF8* and *FGF17*, and no *FGF3* (Supp. Fig. 2f). Comparison with mouse confirmed significant differences from primates, with high expression of *FGF5, 8*, much lower expression of *FGF4,17*, and no expression of *FGF2* in the mouse^21,43,44^.

Of the FGF receptors, only *FGFR1* and *FGFR4* were strongly expressed, with *FGFR1* expression about 5-fold higher (Fig. 2g, Supp. Fig. 2g). Both receptors showed graded expression that decreased from pluripotent to PS-like cells and much lower expression in amnion-like cells (Fig. 2g). Therefore, elevated ERK activity in PS-like cells cannot be accounted for by overall receptor expression. Differential expression of genes regulating heparin-sulfate proteoglycans which modulate FGF receptor binding was mild except for glypicans 3 and 4 which were downregulated in PS-like cells relative to pluripotent cells (Fig. 2h, Supp. Fig. 2g). In contrast, several intracellular negative regulators of ERK signaling were strongly differentially expressed between cell clusters, including *SPRY1* and *PEBP1* which were significantly higher in pluripotent cells (Fig. 2i, Supp. Fig. 2g). Surprisingly, however, expression of *SPRY4* was not elevated in PS-like cells where ERK signaling is high, suggesting that transcription of *SPRY4* may not be a good readout of ERK activity in human hPSCs like it is in the mouse^45^. On the other hand, expression of the ERK negative feedback regulator *IL17RD* (*SEF*) does seem to approximately reflect expected ERK activity in each population. Components of the core ERK pathway including ERK itself were highly expressed across all clusters (Supp. Fig. 2g).

Thus, while scRNA-seq identified specific genes involved, it is consistent with several non-exclusive mechanisms for generating an ERK pattern. For example, uniform FGF2 could form an ERK ring by equally activating receptors across the colony but with ERK response reduced by elevated expression of negative regulators in the pluripotent center. Alternatively, based on gene expression we cannot rule out HSPGs increasing receptor binding specifically in the ring. Finally, localized expression of FGF4 or FGF17 in the PS-like ring could lead to a concentration gradient responsible for increased receptor activation.

### The phosphorylated ERK pattern is due to differential activation of FGF receptors

To distinguish between the different possibilities, we asked whether the spatial pattern of pERK signaling is controlled primarily below, at, or above the level of the receptors. To address this, we first visualized phosphorylated FGFR1 (pFGFR1) and found its spatial pattern strongly resembles that of pERK (Fig. 3a, Supp. Fig. 3a). As expected, pFGFR1 was lost upon brief FGFR inhibition (Fig. 3a, Supp. Fig. 3a). In contrast, after MEK inhibition, high pFGFR1 remained while pERK was greatly reduced (Supp. Fig. 3bc). This suggests the spatial pattern of ERK reflects a spatial pattern of receptor activity.

The spatial pattern in receptor activity could either be due to differential receptor expression or differential receptor activation. There was no significant upregulation of FGF receptors in PS-like cells according to our single cell RNA-sequencing data (Fig 2e). In fact, the most highly expressed receptor, FGFR1, was elevated in pluripotent cells. This suggests the ERK activity pattern is not due to a receptor expression pattern with two caveats. First, mRNA expression may not reflect protein expression. Second, multiple isoforms exist for FGF receptors 1-3 (but not 4) that have different affinity for FGF ligands, leaving open the possibility that PS-like cells could express a different FGFR1 isoform than pluripotent cells, despite overall FGFR1 expression being approximately uniform.

To determine if the spatial pattern in pFGFR1 could be explained by a spatial pattern of FGFR1 protein, we performed immunofluorescence staining of FGFR1. Consistent with the scRNA-seq data, and in sharp contrast to the pFGFR1 stain, FGFR1 protein expression was graded from center to edge with its highest levels in the pluripotent center and therefore could not explain the signaling pattern (Fig. 3b, Supp. Fig. 3d). To determine if different isoforms of FGFR1 are differentially expressed in PS-like vs. pluripotent cells, we designed qPCR primers to detect the IIIb and IIIc isoforms of *FGFR1* (Supp. Fig. 3e-g). These are the most common isoforms, also known as the epithelial and mesenchymal isoforms, and therefore appeared the mostly likely candidates for differential expression. We then analyzed expression of these isoforms in 42h micropatterns versus pluripotent cells and PS-like cells differentiated directly as in Fig. 1d. However, the change in expression between samples was similar for each isoform, with all isoforms decreasing in 2D gastruloids and directly differentiated PS-like cells relative to pluripotent cells (Fig. 3c). To corroborate this and also determine absolute FGFR isoform expression in pluripotent and PS-like cells, we analyzed our single cell RNA-sequencing data and previously published bulk RNA-sequencing data for isoform expression in these two cell types. This revealed that for both pluripotent and PS-like cells FGFR IIIc is the dominant isoform with on average almost 900-fold higher expression than IIIb, while IIIa was not expressed at all (Fig. 3d). Altogether, these data suggest that spatial patterning in ERK is due to differential activation of FGFR receptors, either due to FGF concentration gradients or possibly spatially patterned HSPGs. Immunostaining for HS showed no clear correlation with TBXT or the expected ERK signaling pattern which is similar but extends further inward (Supp. Fig. 4). This does not rule out a role for HSPGs in spatial modulation, since FGF signaling depends on specific sulfation patterns rather than overall HS levels^46,47^. Nevertheless, it led us to focus on FGF gradients as the most likely cause for the ERK signaling pattern.

**Figure 3.**
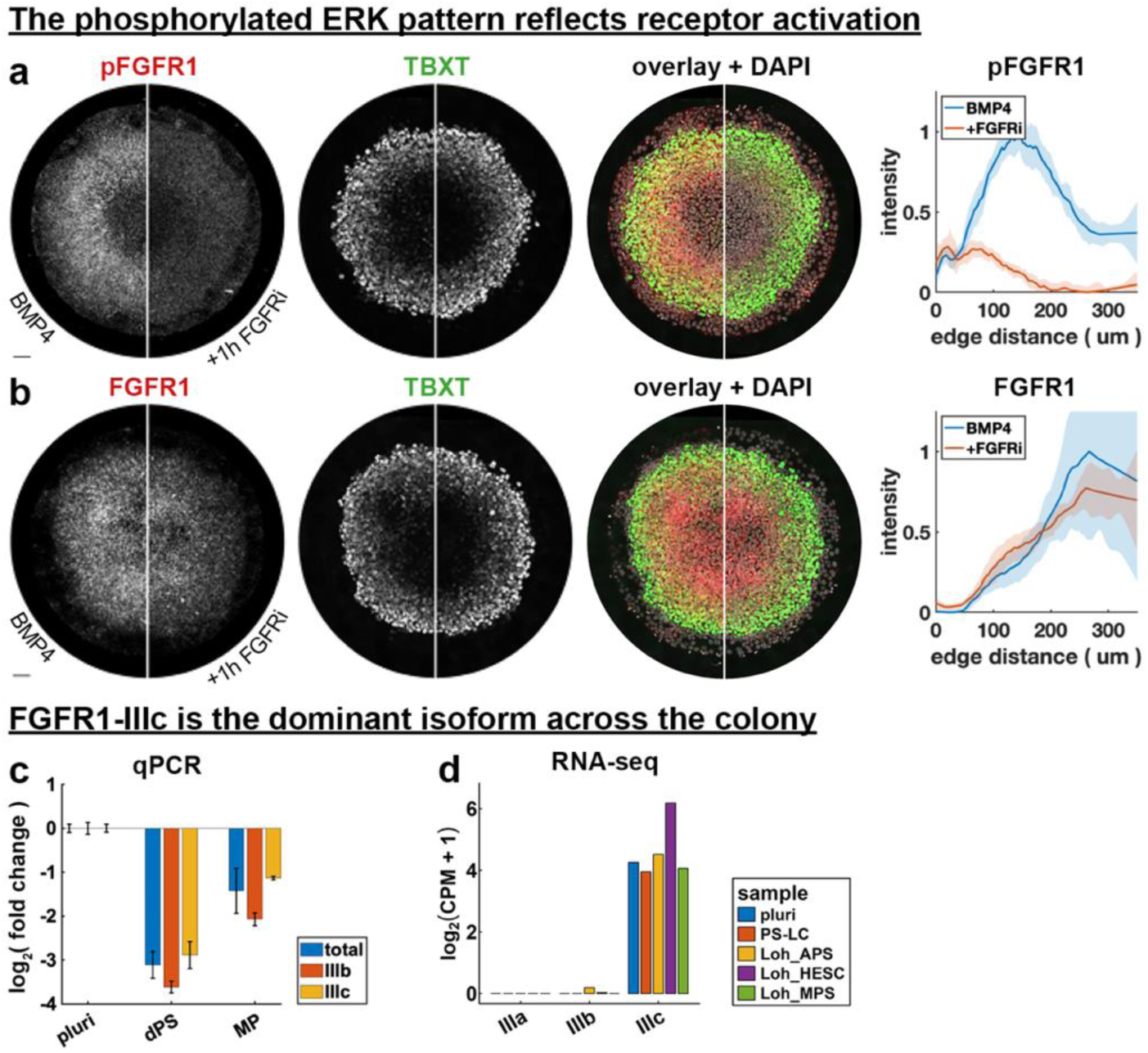
**a)** phospho-FGFR1 stain with and without 1h FGFRi treatment at 41h, radial intensity profile on the right. **b)** FGFR1 with and without 1h FGFRi treatment at 41h, radial intensity profile on the right. Error bars in (b,c) represent standard deviation over N=4 colonies. **c)** qPCR data for relative expression of FGF receptor 1 isoforms IIIb, IIIc and total FGFR1 in pluripotent cells (pluri), gastruloids at 42h (MP), and directly differentiated PS-like cells (dPS). Error bars represent standard deviation of technical triplicates. **d)** Absolute FGFR1 isoform expression from RNA-seq for anterior/mid primitive streak and pluripotent cells from Loh 2016^38^ (Loh_ APS/MPS/HESC) as well as the pluripotent and PS-LC clusters from Fig.2. All scale bars 50 um.

### FGF receptors are localized basolaterally

To maintain an endogenous FGF concentration gradient on the apical side of the colony which faces the medium, would require ligands to remain tethered to the surface. However, FGF has been reported to diffuse freely in the intercellular space^48^. Moreover, there is FGF2 present in the medium that would be expected to uniformly activate FGF receptors. The situation is similar for BMP4, which is present uniformly in the medium for 2D gastruloids but does not cause uniform signaling activity^4^. In the case of BMP, that is because receptors are localized basolaterally^4^ and tight junctions seal the basolateral side from the medium. Consequently, response to exogenous ligands in the medium is restricted to the colony edge, while endogenous ligands are confined to the basal side and prevented from diffusing away into the medium everywhere but the edge. This was found to be critical for patterning in both 2D gastruloids and the mouse embryo^4,5^. To explain the pERK pattern, we therefore hypothesized FGF receptors were similarly restricted to the basolateral side.

To test our hypothesis, we first starved pluripotent colonies of FGF by maintenance in FGF-free E6 medium for 12h and then treated with FGF2. Consistent with our hypothesis, phospho-FGFR and phosphor-ERK were both restricted to the colony edge (Fig. 4a). To determine if this could be explained by diffusion of exogenous FGF from the medium, we repeated the experiment with micropattern pluripotent cells and added fluorescent dextran with a similar molecular weight as FGF. We observed fluorescent dextran diffusing into the intercellular space from the edge inward over the same distance that ERK signaling was observed (Fig. 4b-d). We then acquired high resolution z-stacks of FGFR1 co-stained with the tight junction protein TJP1 (ZO-1) which revealed the majority of FGFR1 was below the tight junctions (Fig. 4e). To directly test that cells are more responsive to basal FGF than apical FGF, we then performed a transwell assay, where pluripotent cells were grown on filters to enable separate apical and basal stimulation. We found that only basal stimulation with FGF led to a significant increase in pERK (Fig. 4f). Finally, to determine if the center of the micropatterned colony is similarly able to respond to basal FGF stimulation and pERK levels there can be attributed to reduced FGF ligand concentration, we performed a scratch assay. Strong ERK response to FGF2 in the media was seen in the colony center around a scratch, but this response was blocked by FGFR inhibitor (Fig. 4gh). If HSPGs rather than FGF ligand were limiting ERK response in the center, we would not have expected strong response around a scratch. The initial ERK levels around the scratch greatly exceeded those anywhere in the control. However, we found this was due to adaptive ERK response to FGF. After six hours ERK levels around the scratch were similar to those in the ERK ring around the PS-like cells (Fig. 4ij). We therefore concluded that the pERK pattern is most likely due to a basolateral FGF ligand gradient which has its maximum in the PS-like ring.

**Figure 4.**
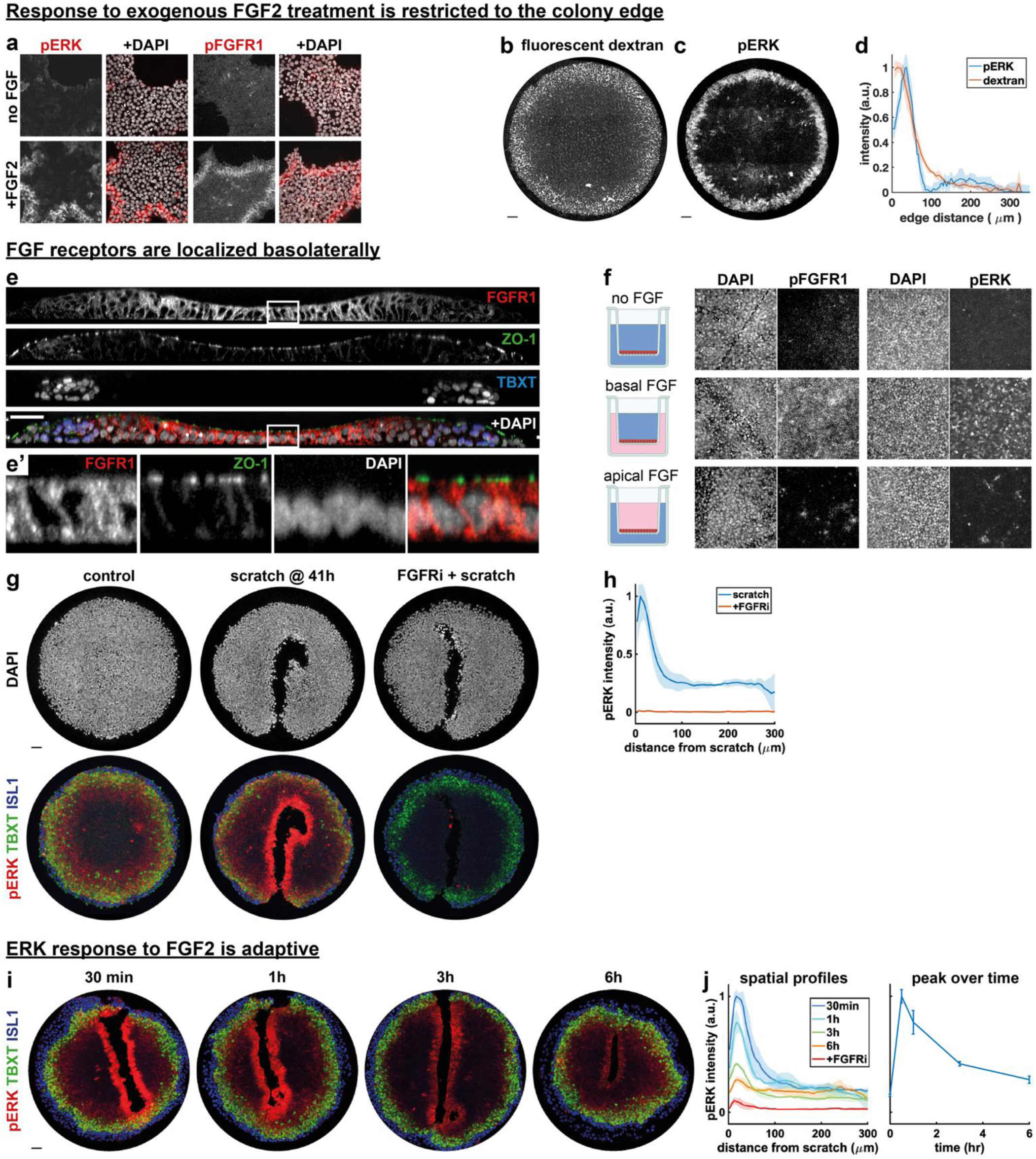
**a)** pERK and pFGFR1 response of pluripotent colonies in standard culture grown in FGF-free (E6) medium for 24 hrs and then treated with FGF2 for 30 minutes. **b,c)** Fluorescent dextran (b) and pERK (c) in micropatterned pluripotent colony 24h after media change containing both FGF2 and dextran. **d)** Radial intensity profile of dextran and pERK for conditions in b,c), averaged over 2 colonies. **e)** Cross-section of 2D gastruloid shows FGF receptor 1 predominantly below the tight junction marked by ZO-1. White box marks are magnified in inset e’. **f)** pERK and pFGFR1 response in transwell experiment with cell treated with FGF either apically or basally. **g)** ERK signaling in scratched colonies with or without FGFRi. **h)** Quantification of pERK intensity as a function of distance from the scratch. Error bars represent standard deviation over colonies. **i)** ERK signaling stains at different times after scratching. **j)** Quantification of pERK intensity as a function of distance from the scratch at different times (left) and peak pERK level over time (right). Error bars represent standard deviation over N=4 colonies. Scale bars 50 micron.

### Endogenous FGF gradients underly the ERK activity pattern

To identify which FGF ligands are required for PS-like differentiation, we first determined the requirement for exogenous FGF2 in the media. Differentiation of 2D gastruloids in FGF-free medium was severely reduced but restored by addition of FGF2 (Fig. 5a, Supp. Fig. 5a). However, if FGF2 was removed 24h after BMP treatment, differentiation was normal. The fact that removal 2h after BMP treatment significantly reduced differentiation suggests this was not a failure to wash out the FGF2. Furthermore, MEK inhibition at 24h severely reduces TBXT expression^34^ while FGF2 is known to have a half-life on the order of hours^49^ and no media changes are performed during standard differentiation. Altogether this suggests that exogenous FGF2 is required at the time of BMP treatment but that endogenous FGFs are sufficient for the ERK activity ring and PS-like differentiation that start a day later.

**Figure 5.**
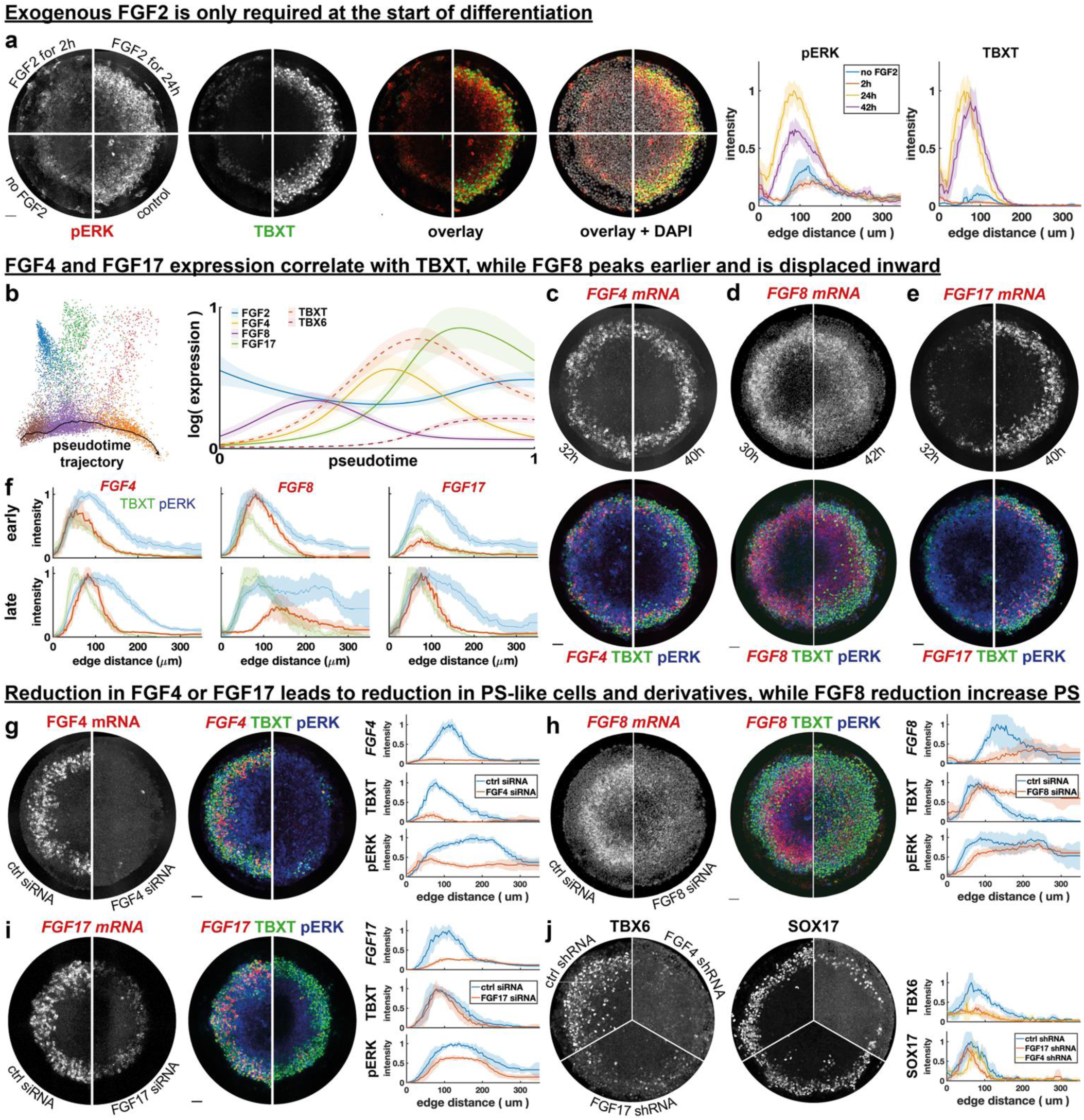
**a)** Pattern at 42h with exogenous FGF2 in the culture medium for different amounts of time. **b)** Pseudotime analysis of FGFs and TBXT, TBX6 expression. **c-e)** FGF4 (c), FGF8 (d), and FGF17 (e) FISH co-stained for TBXT and pERK at 30h or 32h and 40h or 42h after BMP4 treatment. **f)** Radial intensity corresponding to (c-e). **g-i)** FGF4 (g), FGF8 (h), and FGF17 (i) FISH co-stained for TBXT and pERK in control siRNA versus FGF4 siRNA (g), FGF8 siRNA (h) and FGF17 siRNA (i) with corresponding radial intensity profiles. **j)** TBX6 and SOX17 expression in control shRNA versus FGF4 and FGF17 shRNA. Error bars in all panels except (b) represent standard deviation over N=4 colonies. Error bars in (b) represent the 95% confidence interval of the generalized additive model (see methods).

We reasoned that FGFs whose expression correlates with the presence of PS-like cells are more likely to be responsible for the observed ERK ring and PS-like differentiation. Our analysis in figure 2 had implicated FGF4 and FGF17 as most strongly expressed in the PS-like cells and nascent mesoderm, respectively (Fig. 2c). However, it is unclear whether expression of these ligands is a cause or consequence of PS-like differentiation, or both, in the case of a feedback loop. To get more information about the relationship between FGF ligands and PS versus mesoderm differentiation, we performed pseudotime analysis of our scRNA-seq data (Fig. 5b, Supp. Fig 5b). Along a pseudotime trajectory from pluripotent cells to nascent mesoderm, *FGF2* expression was nearly constant. *FGF8* was upregulated at lower levels prior to *TBXT* and declined as *TBXT* was upregulated. *FGF4* upregulation approximately coincided with *TBXT*, but *FGF4* began decreasing as *TBX6* expression increased. In contrast, *FGF17* expression started after *FGF4* but continued to increase after *FGF4* started decreasing. Direct comparison of *FGF* and *TBXT* expression also showed strong positive correlation between *TBXT* and both *FGF4* and *FGF17* but opposing relations with *TBX6* (Supp. Fig. 5c). Together this suggested to us initial TBXT expression may depend more on FGF8 or FGF4, while FGF17 could play a later role in maintaining or expanding the PS-like population or facilitating subsequent mesoderm differentiation.

Pseudotime may not accurately predict true dynamics. Therefore, we verified expression at two different times using RNA FISH (fluorescent in situ hybridization). We found that *FGF4* expression was similar at 32h and 40h (Fig. 5cf, Supp. Fig. 5d). This is consistent with its decrease in pseudotime as cell differentiate to nascent mesoderm, since in real time differentiation is asynchronous and new PS-like cells form while older ones differentiate to nascent mesoderm, which would yield a flatter temporal profile. In contrast, peak *FGF8* levels decreased while *FGF17* strongly increased over time, both matching their pseudotime trends (Fig. 5d-f, Supp. Fig. 5ef). As expected, the spatial profiles of both *FGF4* and *FGF17* resembled TBXT, although the peaks were displaced slightly inward, while *FGF8* expression was highest in the inner part of the ERK ring where TBXT expression was low (Fig. 5c-f). Co-staining showed *FGF4* and *FGF8* expressed in concentric rings with little overlap at 42h (Supp. Fig. 5g). Scatterplots further confirmed that high expression of both *FGF4* and *FGF17* but not *FGF8* was restricted to TBXT positive cells (Supp. Fig. 5h).

To test the functions of these FGFs we decreased their expression. Knockdown (KD) of *FGF4* with small interfering RNAs (siRNA) led to a strong decrease in TBXT and pERK (Fig. 5g, Supp. Fig. 6a). We repeated this in a different cell line and also created stable cell lines expressing short hairpin RNA (shRNA) to knock down *FGF4* in both cell lines with similar effect (Supp. Fig. 6b-d). Strikingly, *FGF8* KD had the opposite effect and led to an increase in TBXT expression without a strong effect on pERK visible at 42h (Fig. 5h, Supp. Fig. 6e). Finally, *FGF17* KD caused only a small decrease in TBXT or pERK, again consistent between different cell lines (Fig. 5i, Supp. Fig. 6f-i). We further interrogated the endoderm and PGC marker SOX17 since both cell types require ERK signaling to differentiate and transiently express TBXT, as well as the mesoderm marker TBX6. TBX6 expression at 42h was near completely lost upon either *FGF4* KD or *FGF17* KD (Fig. 5j, Supp. Fig. 6j). This suggests that while FGF17 is not required for PS induction (Fig. 5i), it is needed for subsequent mesoderm differentiation. In contrast, there was only a small reduction of SOX17 (Fig. 5j). Since we previously found with our protocol most SOX17 positive cells at 42h represent PGC-LCs, while most endoderm differentiates later^34^, this suggests FGF4/17 are not required for PGC-LC induction but does not rule out a requirement for these FGFs in endoderm differentiation. A double KD of *FGF4* and *FGF17* had a similar phenotype as *FGF4* KD alone, suggesting FGF17 may be downstream of FGF4 (Supp. Fig. 6k). Combined, these data support distinct functions for FGF4, FGF8, and FGF17 in our human gastrulation model, where FGF4 and FGF8 may play opposing roles in PS induction while FGF17 is important during continued differentiation to mesoderm.

## Discussion

We have shown that in a stem cell model for early human gastrulation, FGF-dependent ERK signaling is directly required for differentiation to PS-like cells and their derivatives, and that loss of differentiation upon FGF inhibition can be rescued by directly activating ERK. We also showed that FGF receptors are polarized basolaterally, preventing the colony center from responding to exogenous FGF in the medium. Furthermore, FGF receptor activation is increased in the PS-like ring, matching the pattern of active ERK despite overall receptor expression being lower in the PS-like ring than in the pluripotent center. This suggests elevated ERK activity is due to increased binding of FGFs to the receptors in this region, possibly due to increased FGF concentration. We found that both FGF4 and FGF17 are highly expressed in PS-like cells, while FGF8 is expressed earlier and displaced inward (Fig.6). Knockdown of FGF4 leads to a strong reduction in ERK activity and PS-like cells, suggesting it may be responsible for increased ERK signaling in the PS-like region. In contrast, FGF8 knockdown led to an expansion of ERK activity and PS-like differentiation. Finally, FGF17 knockdown leds to a loss of mesoderm differentiation.

**Figure 6.**
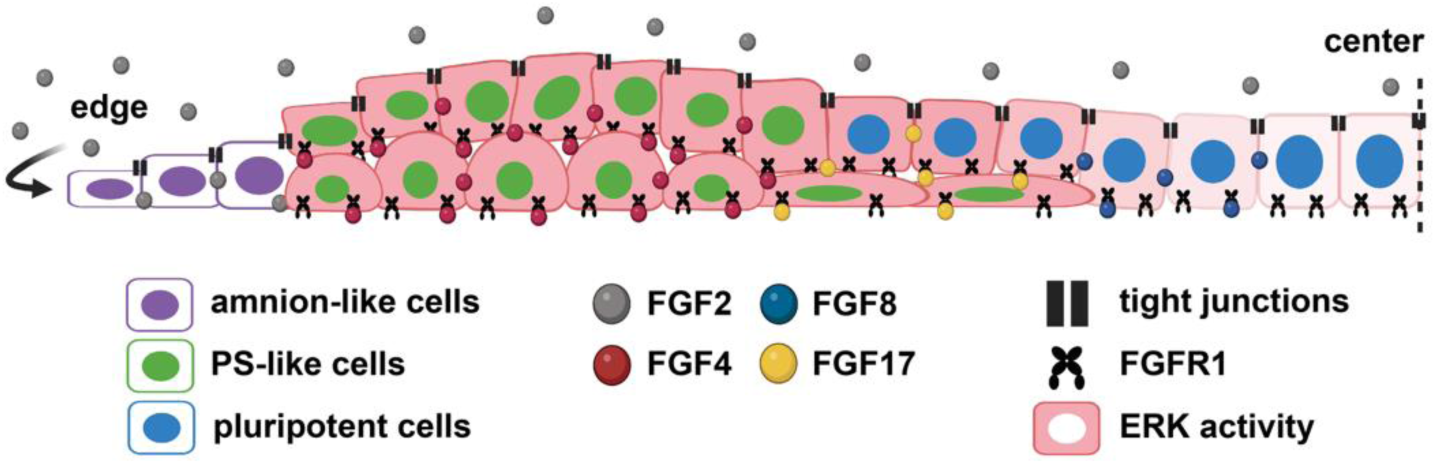
Graphical summary. Exogenous FGF2 acts on the colony edge and is prevented from reaching the basal FGFR1 receptors elsewhere by tight junctions. Primitive streak cells express FGF4 and FGF17 and these FGFs are associated with high ERK activity. FGF8 is expressed further inside the colony.

Many questions remain. Although we showed that FGF gradients are likely the dominant cause for the spatial pattern of ERK signaling, our scRNA-seq data suggest other levels of regulation may contribute, which remain to be explored. Furthermore, although we showed FGF/ERK signaling is directly required for PS-like differentiation, it may also act indirectly by affecting the expression of Wnt and Nodal. We will explore this and the more general question of how FGF integrates with the hierarchy of BMP, Wnt, and Nodal in a separate manuscript. Related is the indirect effect FGF may have on patterning by its control of cell growth and thereby cell density. Cell density affects the response to exogenous BMP and downstream expression of Wnt, Nodal, and various inhibitors^4,50,51^, as well as FGF itself, possibly causing a “community effect”^52–54^. It likely has other effects as well. However, the rescue of TBXT expression but not growth by ERK activation when FGF receptors are blocked suggests the FGF-dependent cell growth is not required to generate a primitive streak-like ring. The ability of exogenous Wnt and Nodal to induce PS-LCs in very sparse culture while MEK and FGFR inhibition block differentiation at the same density similarly argues against a requirement for FGF-dependent cell growth in PS-LC differentiation (Supp. Fig. 1h).

Another important question is to what extent different FGFs act redundantly or perform different functions. Our data suggest a picture where different FGFs are required consecutively, starting with exogenous FGF2, then endogenous FGF4, to support initial differentiation of PS-like cells, and finally FGF17 to maintain and expand the PS-like ring and support mesoderm and possibly endoderm differentiation. Whether these FGFs could substitute for each other if they were expressed at similar times and levels is unclear. It also remains to be determined whether the nearly constant high expression of FGF2 we observed is required for patterning. Most puzzling is the role of FGF8, whose knockdown led to an increase in PS-like differentiation. Given that FGF8 expression is displaced inward, one possibility is that it induces inhibitors of other PS-inducing signals there in order to limit PS differentiation, e.g., the Nodal inhibitor Lefty, which is expressed in a similar domain^55^. The rescue of PS-like differentiation by uniform activation of ERK to a supraphysiological level suggests that the role of the different FGFs is simply to sufficiently activate ERK signaling. However, future work will have to investigate the dox-SOS^cat^ rescue phenotype in much greater detail to see how differentiation and other cell behaviors such as migration are affected.

It is interesting to ask if there is a simple explanation for the seemingly large differences in FGF expression between primates and other species. FGF8 is the only FGF shown to be required for mouse gastrulation and is upstream of FGF4 in both chick and mouse, but it is barely expressed in human and monkey primitive streak and appears to play a different role in our stem cell model. If FGFs are interchangeable in simply activating ERK for primitive streak induction, variation in FGF expression is not surprising. However, FGF4 and FGF8 are the main FGFs in mouse, chick and frog gastrulation^56–58^, suggesting a high degree of conservation. It is possible FGF8 has simply been substituted by FGF17, which is in the same subfamily^9^, but this does not explain why the role of FGF8 in gastrulation otherwise appears strongly conserved, or why FGF8 is upstream of FGF4 in the mouse while FGF4 is expressed before FGF17 in 2D gastruloids. Similarly, it is possible that FGF2 in primates substitutes for FGF5 in the mouse as the FGF that is highly expressed in the epiblast. Although these are not in the same subfamily and generally have different receptor affinities, they both have highest affinity for FGFR1^9^. That leaves FGF4 as the only FGF that may be conserved across vertebrate gastrulation, and it is tempting to speculate on why. In Xenopus, FGF4 and TBXT (Xbra) function in a positive feedback loop^22,59^. The strongly correlated initial transcriptional dynamics of TBXT and FGF4 in 2D gastruloids are consistent with a similar feedback loop. However, an FGF8 feedback loop has also been proposed and recent work suggests TBXT is not required for FGF4 or FGF17 expression^54,60,61^. Future work will therefore have to investigate further if FGF4 plays a unique role.

A final related question is the origin of the discrepancies we and others^36^ found between the primate (i.e., non-mouse) samples in Fig. 2 and Supp. Fig. 2: FGF3 appears only significantly expressed in the human embryo, but not in the monkey embryo and human stem cell model. This could also reflect genetic variation: if FGFs function redundantly, their relative expression levels could be relatively unconstrained between individuals. On the other hand, in the nascent mesoderm, FGF2 is only high in human gastruloids. This may reflect developmental stage: the in vivo data is developmentally more advanced and nascent mesoderm may lose FGF2 expression later. It is also possible the maintenance conditions of hPSCs lead to in vitro artefacts in FGF expression.

In conclusion, the complexity of FGF signaling, with its many ligands, modulators, and functions, has prevented a clear understanding of its role in mammalian gastrulation in general and in human gastrulation specifically. Here we showed that 2D human gastruloids provide a powerful approach to make headway and although many open questions remain, we have taken significant steps towards this goal.

## Acknowledgements

We thank Lila Solnica-Krezel, Blerta Stringa, Aryeh Warmflash, and Ben Allen for discussions. We thank Jared Toettcher for the dox-SOScat plasmid. Library prep and next-generation sequencing was carried out in the Advanced Genomics Core at the University of Michigan. This work was supported by the National Institute of General Medical Sciences (NIGMS R35GM138346), and the Branco Weiss Fellowship – Society in Science. KJ was partially supported by the Michigan Pioneer Postdoctoral Fellowship and the NIH F32 Ruth L. Kirschstein Postdoctoral National Research Service Award (5F32HD108980-02).

## Data availability

Raw single cell RNA sequencing data generated in this study was deposited in GEO (GSE271604), analyzed data will be added upon publication. Raw image data are available upon request.

## Code availability

All code for data analysis and model simulations is available on https://github.com/idse/FGF

## Methods

### Cell lines

The cell lines used were the embryonic stem cell line ESI017 (XX), and the induced pluripotent stem cell line PGP1 (XY). The pluripotency of these cells was confirmed by immunostaining of pluripotency markers OCT3/4, SOX2, NANOG. All cells were routinely tested for mycoplasma contamination, and negative results were recorded. ESI017 cells stably expressing doxycycline inducible SOS^cat^ were made using a published Piggybac plasmid gifted by Jared Toettcher^37^.

### Cell culture and differentiation

Human pluripotent stem cells were cultured in the commercially defined pluripotency maintaining media mTeSR1 (STEMCELL Technologies #85850) on Cultrex (R&D Systems)-coated tissue culture plates. For FGF2 starvation we used Essential 6 Medium (Thermofisher, A1516401), or mTeSR1 without bFGF, TGF-beta, LiCl, GABA and pipecolic acid (STEMCELL Technologies #05896). For routine cell maintenance, cell passaging was performed every 3 days. For whole-colony passaging, L7^62^ was used, while single-cell passaging was performed with Accutase or TrypLE (Gibco). Single-cell suspensions were used to seed for all the differentiation experiments to control initial cell number.

To directly differentiate to primitive streak-like cells, cells were resuspended in a single-cell suspension and seeded in cultrex coated wells in mTeSR with ROCK inhibitor (RI) Y-27632 (MeChemExpress, cat# HY-10583). Cells were then maintained in mTeSR with RI for 24h before adding CHIR (3-5uM) and Activin (100ng/ml) treatment for 24-30h. Experiments were performed in 18-well Ibidi slides (cat# 81818).

To differentiate cells for 2D gastruloids by micropatterning, we followed the protocol in^42^. Briefly, cells were resuspended in a single-cell suspension and seeded in laminin-coated micropatterned wells in mTeSR with ROCK inhibitor (RI) (MeChemExpress, cat# HY-10583). Micropatterned wells were washed with PBS-/- 30 minutes after the initial seeding to wash off the non-attached cells. Cells were then maintained for 24 hours in mTeSR before adding the BMP4 treatment by a full medium change. For time series micropatterned differentiation, full media changes with designated treatment(s) were performed every 24 hours. Experiments were performed in micropatterned 18-well Ibidi slides (cat# 81818) prepared as previously described^63^. Cell signaling reagents and doses used are listed in Supplementary Table 1.

### Immunofluorescence staining

Antibodies can be found in Supplementary Tables 2 and 3. For stains including pERK, we used methanol fixation, for all other stains we used PFA fixation. PFA fixation: Samples from the 18-well Ibidi slides were rinsed with PBS, fixed for 20 min in 4% paraformaldehyde, rinsed twice with PBS, and blocked for 30 min at room temperature with 3% donkey serum and 0.1% Triton X-100 in 1× PBS. After blocking, cells were incubated with primary antibodies at 4°C overnight, followed by three washes in PBST (PBS with 0.1% Tween 20). They were then incubated with secondary antibodies and DAPI for 30 min at room temperature and washed twice in PBST at room temperature. Methanol Fixation: Samples from the 18-well Ibidi slides were rinsed with PBS, fixed for 20 mins in cold methanol, then rinsed with PBS for 5 min two times, after which 10% phosphate buffered formalin was added for 20 min at room temperature. Cells were then rinsed with TBS twice followed by 5 mins of incubation in cold methanol and rehydrate in TBS for 20 mins. Blocking was done for 30 mins at room temperature with 5% donkey serum, 1% BSA, 0.2% Triton-X-100 in 1x TBS after which staining proceeded the same way as with PFA fixation.

### Microscopy

Fixed sample imaging was performed with an Andor Dragonfly/Leica DMI8 spinning disk confocal microscope with a 40x, 1.1NA water and 20x 0.8NA air objectives using Andor Fusion software version 2.3.0.31 and a Nikon/Yokogawa spinning dish confocal microscope with a 40x silicon oil objective using NIS Elements AR software version 5.41.02. Live-cell imaging was performed with an Andor Dragonfly/Leica DMI8 spinning dish confocal microscope under the controlled temperature (37°C), CO_2_ concentration (5%), and humidity (>60%).

### General image analysis

We segmented nuclei in individual z-slices based on nuclear fluorescence stained with DAPI in fixed cells using a pipeline we previously described, which integrates two machine learning approaches: Ilastik pixel classification^64^ and Cellpose^65^ (v1). Fluorescence intensities were then calculated per nucleus as mean intensities in the 3D nuclear again with our established custom image-processing pipeline. To calculate the radial fluorescence distributions in micropatterned colonies, colonies were subdivided into radial bins with equal numbers of cells, mean or median and variance were then calculated within these bins. To calculate positive fractions, markers were first thresholded based on their intensity distribution, after which the fraction of cells positive for each marker was calculated within different bins. Standard deviations were then calculated over multiple replicate colonies. For visualization purposes only, background subtraction using grayscale opening on a 100 micron scale and 5 pixel wide median filter was applied slice by slice to z-stacks of micropatterned colonies before making maximal intensity projections.

### (Single-cell) RNA sequencing and analysis

Cells were collected using accutase and resuspended in ice-cold PBS. Single-cell RNA-sequencing was performed by the University of Michigan Advanced Genomics Core. Cells were barcoded using the 10X Genomics Chromium system (part numbers 1000268, 1000120, 1000215). For quality control, cDNA was quantified by Qubit High Sensitivity DNA assay and Agilent TapeStation. Sequencing was performed on the Illumina NovaSeq 6000 with NovaSeq S4 flowcell and Control Software version 1.7.0. Reads were aligned using cellranger-4.0.0 with the GRCh38 reference. Further processing was mostly done in Python, primarily using the Scanpy^66^ and SCVI^67^ packages. Analysis was performed on log transformed data. Library size normalization was performed on the intersection of the genes present in our data and the reference CS7 human embryo data to facilitate comparison of expression levels. We integrated the two scRNA-seq datasets for 42h 2D human gastruloids using SCVI based on the top 2000 highly variable genes, which were selected using the default function in Scanpy using the seurat_v3 flavor. Cell cycle genes were regressed out during integration. Dimensional reduction for visualization was performed using PHATE^39^ on the SCVI latent space. Gene expression was smoothened with MAGIC^68^ for visualization on PHATE plots in Fig. 2e,g-i. To annotate different cell types, we used MAGIC to impute that data and get smooth distributions of gene expression. We then thresholded marker genes to assign cell fate (Table 4), which produced better results than the more common unsupervised clustering using Leiden, and in particular more consistent results between datasets from different species. For interspecies comparison of FGF ligands we normalized the to the maximal FGF expression within each species, where for the mouse data, the three different time points were first integrated using SCVI. Thus we compare between species the relative expression of FGF ligands within each species, e.g., FGF8 expression is much higher than FGF4 at any time in the mouse, but much lower than FGF4 in human.

For pseudotime analysis (Fig. 5), we applied the trajectory inference tool Slingshot^69^ in R to compute the pseudotime trajectory in the SCVI latent representation. Slingshot determines the membership of a cell to a lineage at each branching point by its projection distance to the fitted principal curve of that lineage. We projected the mesoderm developmental trajectory onto the PHATE map for visualization. We further used TradeSeq^70^, which fits a generalized additive model (GAM), to infer the pseudotime gene expression trends along the mesoderm lineage trajectory from Slingshot. For visualizing the correlation between the expression of FGF ligands and marker genes (Supp. Fig. 5), we used our SCVI model to denoise the expression and mutual nearest neighbor to further harmonize the difference between samples.

To quantify isoforms from RNA sequencing salmon v1.10.0 was used^71^. First, gencode.v45.transcripts.fa was downloaded from gencodegenes.org. The reference was customized by removing lines for FGFR1 and replacing them with the transcript sequence for FGFRIIIa, FGFRIIIb, FGFRIIIc (sequences included as supplementary file). Three samples from GEO dataset GSM2257301 were downloaded through SRA (SRX1725562 (Pluripotent), SRX1725577 (APS), SRX1725643 (MPS))^38^. Adapters and the first 10 bases were trimmed from the FASTQ files using trim_galore. Finally, the Salmon “quant” command with the --validateMappings option was used to quantify transcripts from trimmed FASTQ files. For single cell datasets, https://timoast.github.io/sinto/ was used to subset the 10x aligned file by cluster. Reads for clusters of interest were converted back to raw FASTQ files and run through the same alignment-based mode of salmon but as single end reads.

### Quantitative real-time PCR

RNA were extracted using RNAqueous™-Micro Total RNA Isolation Kit (CAT# AM1931, Thermo Fisher Scientific) and then cDNA synthesis was prepared from it using SuperScript™ VILO™ cDNA Synthesis Kit (CAT# 11754250, Thermo Fisher Scientific) following the manufacturer‘s protocol with adjustment for optimization. Measurements were performed with SYBR green and the primers in the Table 5. GAPDH was used for normalization in all experiments.

### Fluorescent dextran assays

Fluorescent dextran essays were performed by adding 10 ug/ml fluorescein isothiocyanate-dextran (10KD, Sigma Aldrich) to the media. To calculate fluorescence intensity as a function of distance from the colony edge, Otsu thresholding followed by several morphological operations was used to create a mask for the colony in each z-slice, thereby excluding bright dextran fluorescence outside the colony. Background subtraction on each z-slice of the image was performed to further remove background signal from dextran outside the colony while maintaining signal in the intercellular space. The radial intensity profile was then calculated from the maximal intensity projection of the masked, background subtracted image.

### Scratch assays

Human embryonic or induced pluripotent stem cells were differentiated on micropattern wells for 41h, then each micropattern colony was manually scratched with a serological needle. After 60mins, they were fixed with PFA or methanol followed by immunostaining with appropriate antibodies. To calculate the intensity profile as a function of distance from a scratch, we created a mask for the scratch based on the difference between an overall mask for the colony and the convex hull, calculated the distance transform of the scratch mask, then binned pixels by distance from the scratch and calculated their mean ERK intensity.

### Transmembrane assays

Human embryonic or induced pluripotent stem cells were seeded into Corning 6.5 mm transwell with 0.4 µm pore polyester membrane insert (cat# 3470) with E6 supplemented with TGF1b, FGF2, and Y-27632 for 16h, then media was changed to E6 with TGF1b. After 24h, the media was changed to E6 supplemented with FGF2 from the top or the bottom side of the transwell. 30 mins later, cells were fixed followed by immunofluorescence staining. The transwell cartoon in Fig 4 was generated with Biorender.

### Fluorescence In Situ Hybridization

The fluorescence in situ hybridization protocol (FISH) was performed according to the manufacturer’s instructions (ACDbio, RNAscope multiplex fluorescent manual). A list of probes and reagents can be found in Supplementary Table 6.

### Gene targeting by shRNA

We used the constructs shRNA F4: pPB[shRNA]-EGFP-U6>hFGF4[shRNA#1] and shRNA F17: pPB[3shRNA]-U6>hFGF17[shRNA#1]-U6>hFGF17[shRNA#2]-U6>{hFGF17_shRNA3}-hPGK>Bsd both purchased from VectorBuilder in a PiggyBac Transposon Vectors with puromycin resistance which was stably integrated into ESI17 hESCs and PGP1 hiPSCs. Transfections were performed using Lipofectamine™ Stem Transfection Reagent (CAT# STEM00001, Thermo Fisher Scientific), following the manufacturer‘s protocol.

### Gene targeting by siRNA

We purchased predesigned siRNA from Millipore Sigma; NM_002007 (FGF4), NM_006119 (FGF8), NM_008004 (FGF17). The siRNA transfections were performed using Lipofectamine™ RNAiMAX Transfection Reagent (CAT# 13778075, Thermo Fisher Scientific), following the manufacturer‘s protocol with adjustment for optimization.

### Statistics and reproducibility

All experiments were performed twice or more. All the attempts in replicating experiments yielded consistent results. Unless specifically noted, all the quantifications were performed with N=4 colonies within the same experimental condition.

### Ethics statement

We comply with all the ethical regulations in our research. We were granted approval by the Human Pluripotent Stem Cell Research Oversight (HPSCRO) Committee at the University of Michigan to work with human embryonic stem cells and human induced pluripotent stem cells.

---

**Supplementary Table 1:**
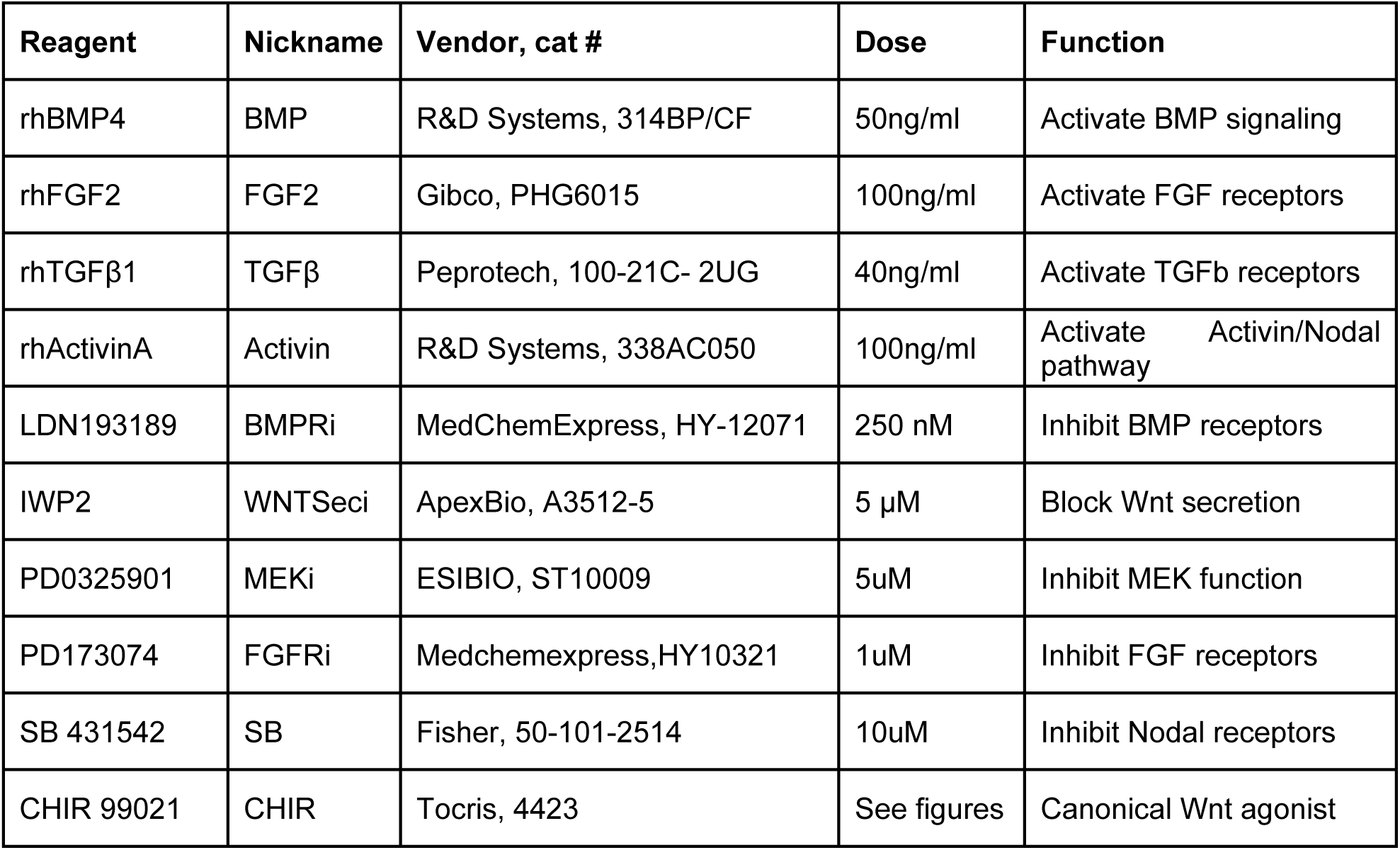
Cell signaling reagents.

**Supplementary Table 2:** Primary antibodies used for immunofluorescence.

| Antigen | Species | Dilution | Cat # | Vendor |
| --- | --- | --- | --- | --- |
| ISL1 | Mouse | 1:200 | 39.4D5 | DSHB |
| SOX2 | Rabbit | 1:200 | 3579S | Cell Signaling Tech |
| TBXT (BRA) | Goat | 1:300 | AF2085 | R&D Systems |
| EOMES | Mouse | 1:200 | MAB6166 | R&D Systems |
| MIXL1 | Rabbit | 1:150 | A17223 | ABclonal |
| pERK | Rabbit | 1:200 | 9101S | Cell signaling Tech |
| pFGFR1 | Rabbit | 1:100 | 52928 | Cell signaling Tech |
| FGFR1 | Rabbit | 1:100 | 9740T | Cell signaling Tech |
| Heparan Sulfate | Mouse | 1:100 | 370255-S | AMS Bio |

**Supplementary Table 3:** Secondary antibodies.

| Antigen | Species | Dilution | Cat # | Vendor |
| --- | --- | --- | --- | --- |
| Alexa Fluor 488 anti-mouse | Donkey IgG | 1:500 | A21202 | ThermoFisher Scientific |
| Alexa Fluor 555 anti-rabbit | Donkey IgG | 1:500 | A31572 | ThermoFisher Scientific |
| Alexa Fluor 647 anti-goat | Donkey IgG | 1:500 | A21447 | ThermoFisher Scientific |
| Alexa Fluor 647 anti-rabbit | Donkey IgG | 1:500 | A31573 | ThermoFisher Scientific |
| Alexa Fluor 555 anti-goat | Donkey IgG | 1:500 | A21432 | ThermoFisher Scientific |

**Supplementary Table 4:** Marker genes for cell fate annotation.

| Cell type | Markers |
| --- | --- |
| Pluripotent | SOX2+POU5F1+TBXT-MIXL1-SOX17- |
| Amnion-like cell (AM-LC) | ISL1+POU5F1+TBXT-MIXL1-SOX17- |
| Primitive streak-like cell (PS-LC) | TBXT+MIXL1+POU5F1+TBX6-SOX17-FOXA2- |
| Nascent mesoderm (NasMe) | TBXT+MIXL1+TBX6+SOX17- |
| Endoderm | SOX17+EOMES+TFAP2C- |
| Ectoderm | SOX2+POU5F1-TBXT-MIXL1-SOX17- |
| Primordial germ cell-like cells (PGC-LC) | SOX17+TFAP2C+ |

**Supplementary Table 5:**
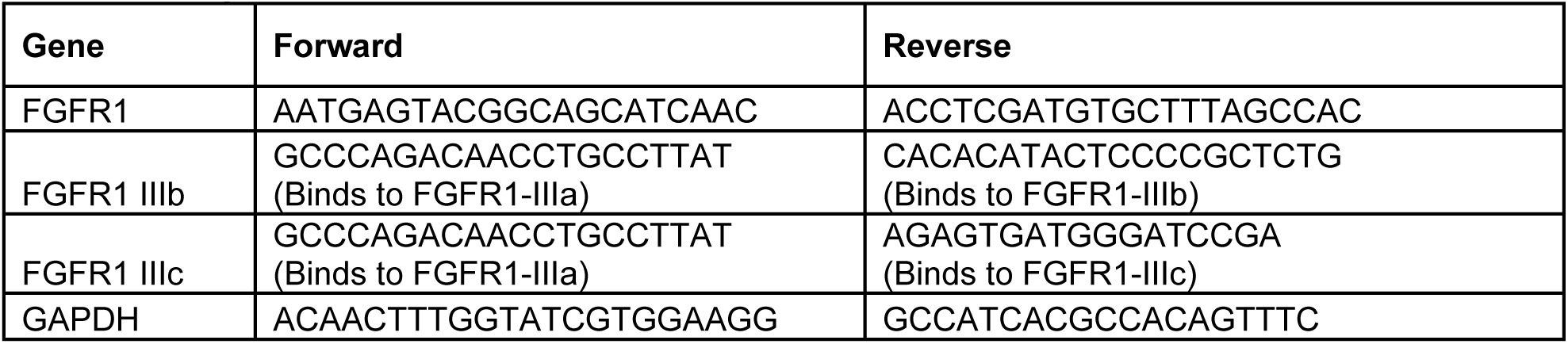
qPCR primers.

**Supplementary Table 6:** Probes for Fluorescent In Situ Hybridization.

| Probe | TSA dilution | Cat # | Vendor |
| --- | --- | --- | --- |
| Hs-FGF4 | 1:2000 | RNAScope #512291 | ACDBio |
| Hs-FGF8-C2 | 1:2000 | RNAscope 415791-C2 | ACDBio |
| Hs-FGF17-C1 | 1:2000 | RNAscope #1148351-C1 | ACDBio |

## Supplementary Figures

**Supplementary Figure 1.**
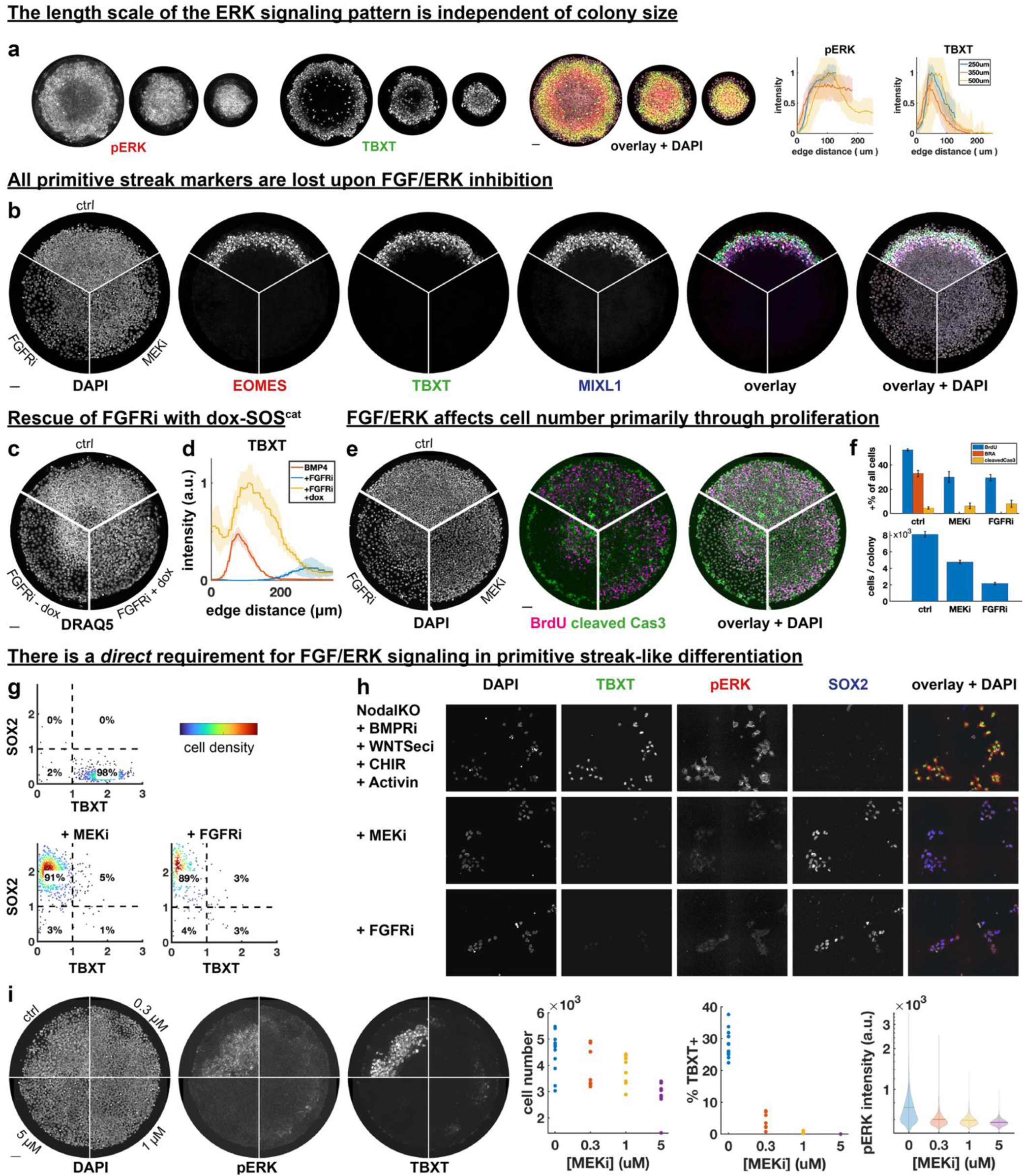
**a**) Stains and quantification for pERK and TBXT in smaller colonies, diameter from left to right: 500um, 350um, 250um. **b)** Staining for three primitive streak markers with or with FGFRi or MEKi. **c)** DAPI stain corresponding to Fig. 1d. **d)** Radial intensity profile for TBXT corresponding to Fig. 1d. **e**) BrdU and cleaved Cas3 stainings after MEK or FGFR inhibition. **f)** Quantification of conditions in e. **g)** Quantification of conditions in Fig. 1g for five images each, N indicates total number of cells. **h**) Low density differentiation with and without MEK and FGFR inhibition. **i)** pERK and TBXT stains and quantification for different doses of MEK inhibitor shows low doses effectively block differentiation with minimal impact on cell number.

**Supplementary Figure 2.**
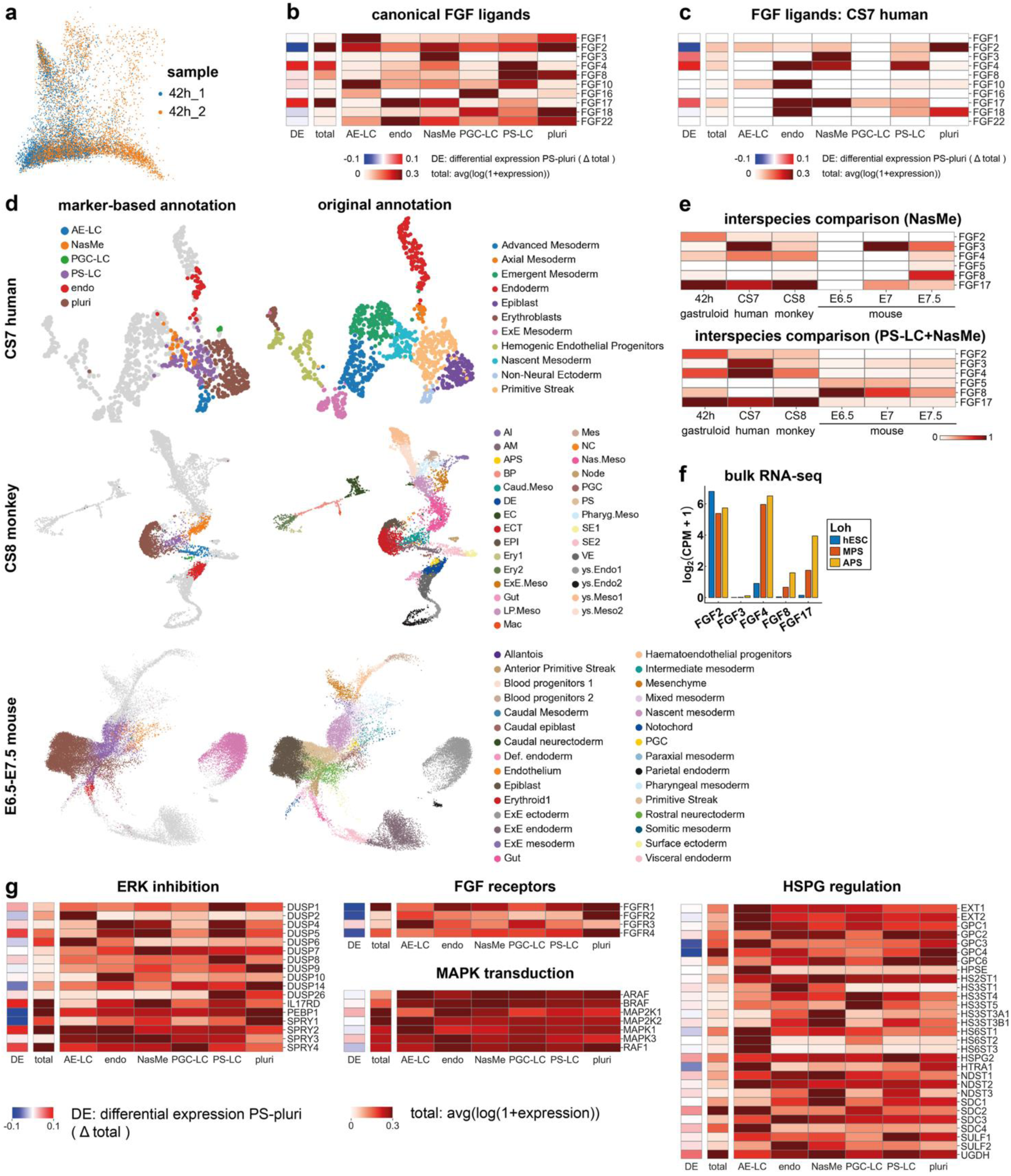
**a)** PHATE projection of scRNA-seq data colored for sample. **b,c)** Expression of canonical FGF ligands across different cell types in 2D gastruloids (b) and human CS7 embryo (c). Expression across clusters is normalized by row, while total expression is shown in a separate column to the directly to left. The leftmost column indicates differential expression between PS-like and pluripotent cells. The color scale of (differential) expression is cut off for better contrast. **d)** Threshold-based annotation (left) for human, monkey, and mouse embryos used in interspecies comparison (Fig. 2f), compared to original annotation (right). **e)** Interspecies comparison as in Fig. 2f but for nascent mesoderm and PS-LC + nascent mesoderm. **f)** Bulk RNA-seq data by Loh et al^38^ for expression of different FGFs in directed differentiation to mid primitive streak (MPS) or anterior primitive streak (APS) relative the human embryonic stem cells (hESCs). **g)** Expression of genes involved in FGF signaling.

**Supplementary Figure 3.**
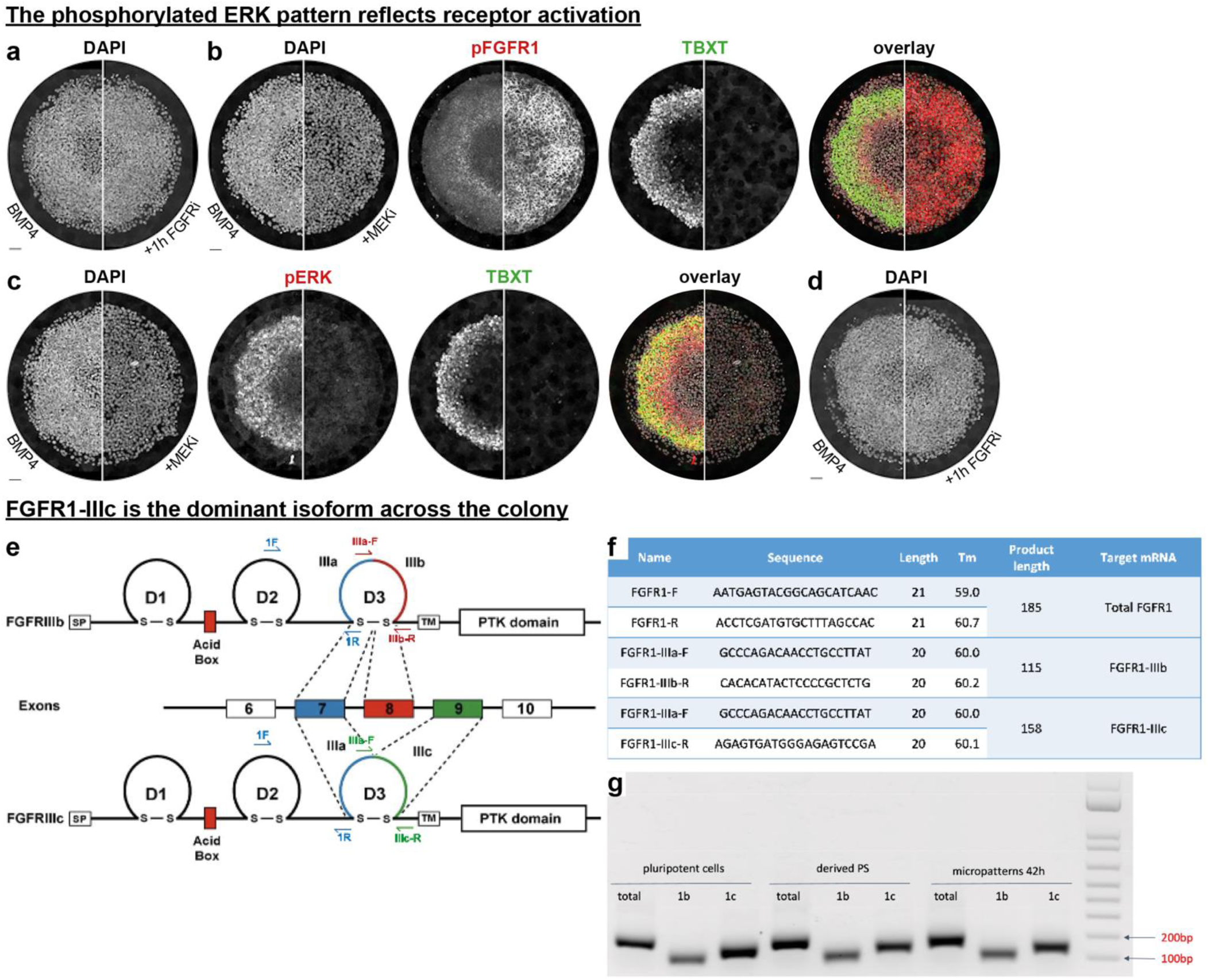
**a)** DAPI image corresponding to Fig. 3a. **b-c)** pFGFR1 (b) or pERK (c) and TBXT stains with and without continuous MEKi treatment. **d)** DAPI image corresponding to Fig. 3b. **e)** Design of qPCR primers to detect FGFR1 isoforms, diagram adapted from Eswarakumar et al^72^. **f)** Primer sequence and properties. **g)** Amplicon sizes match prediction. Scale bars 50um.

**Supplementary Figure 4.**
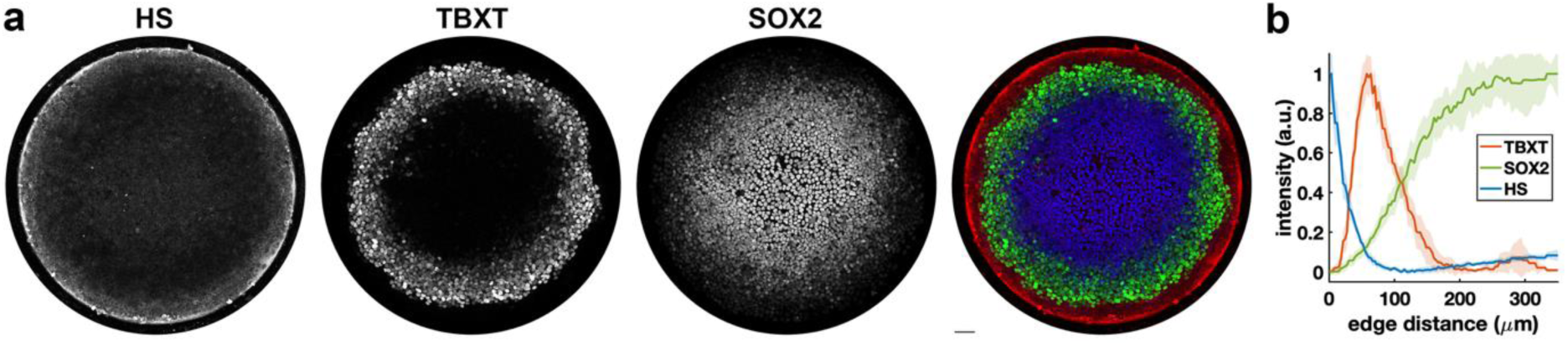
**a,b)** Heparan sulfate (HS) stain (a) and radial intensity profile (b). Scale bar 50um. Error bands represent standard deviation over N=4 colonies.

**Supplementary Figure 5.**
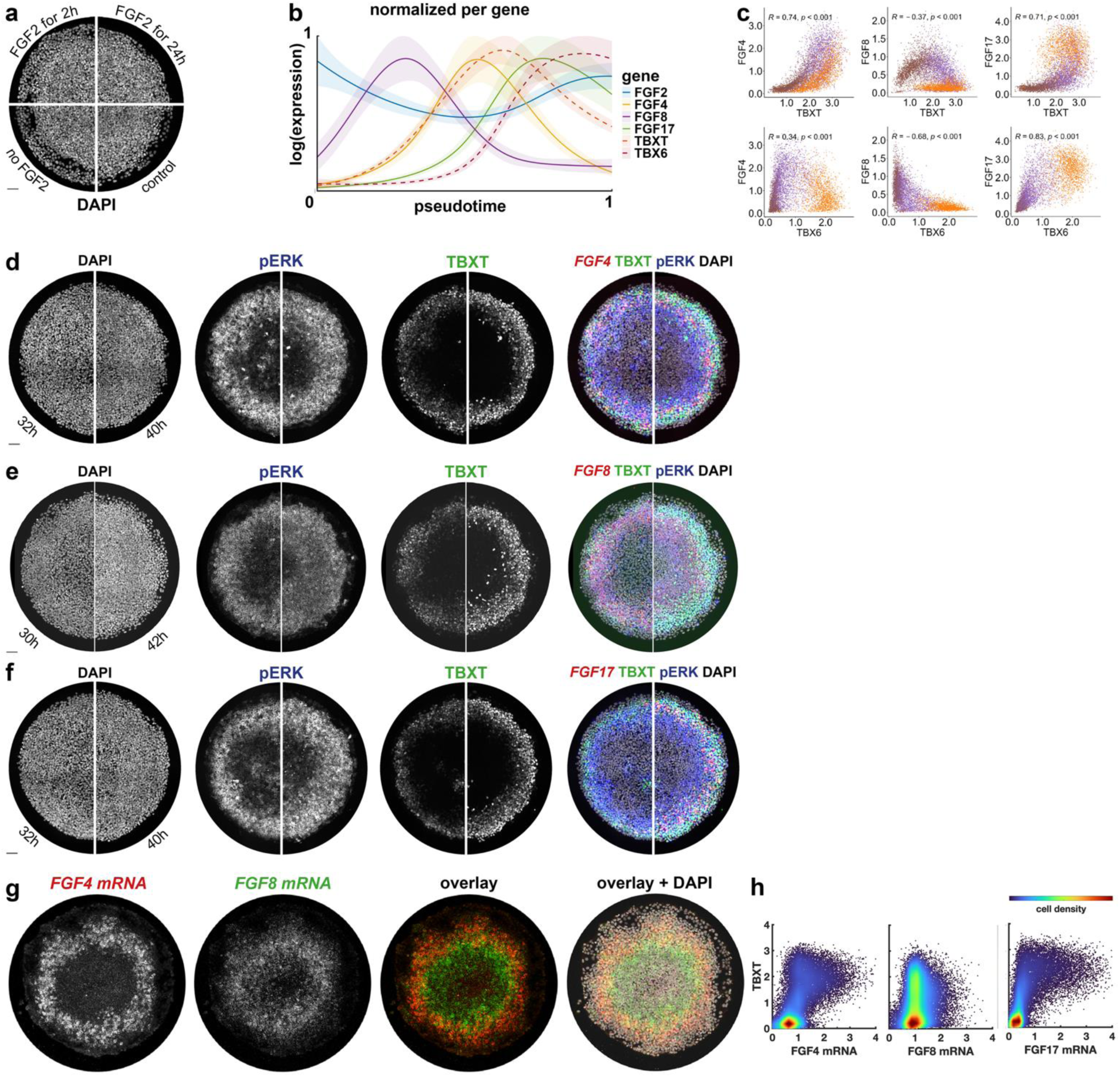
**a)** DAPI image corresponding to Fig. 5a. **b)** Pseudotime dynamics from Fig. 5b with each gene normalized individually. **c)** Correlations between FGFs and TBXT, TBX6 colored for clusters, with cluster colors matching Fig. 2d. **d-f)** DAPI images and overlays including DAPI corresponding to Fig. 5c-e, respectively. **g)** Combined FISH for FGF4 and FGF8. **h)** Scatterplots of FGF4, FGF8, and FGF17 mRNA versus TBXT colored for cell density.

**Supplementary Figure 6.**
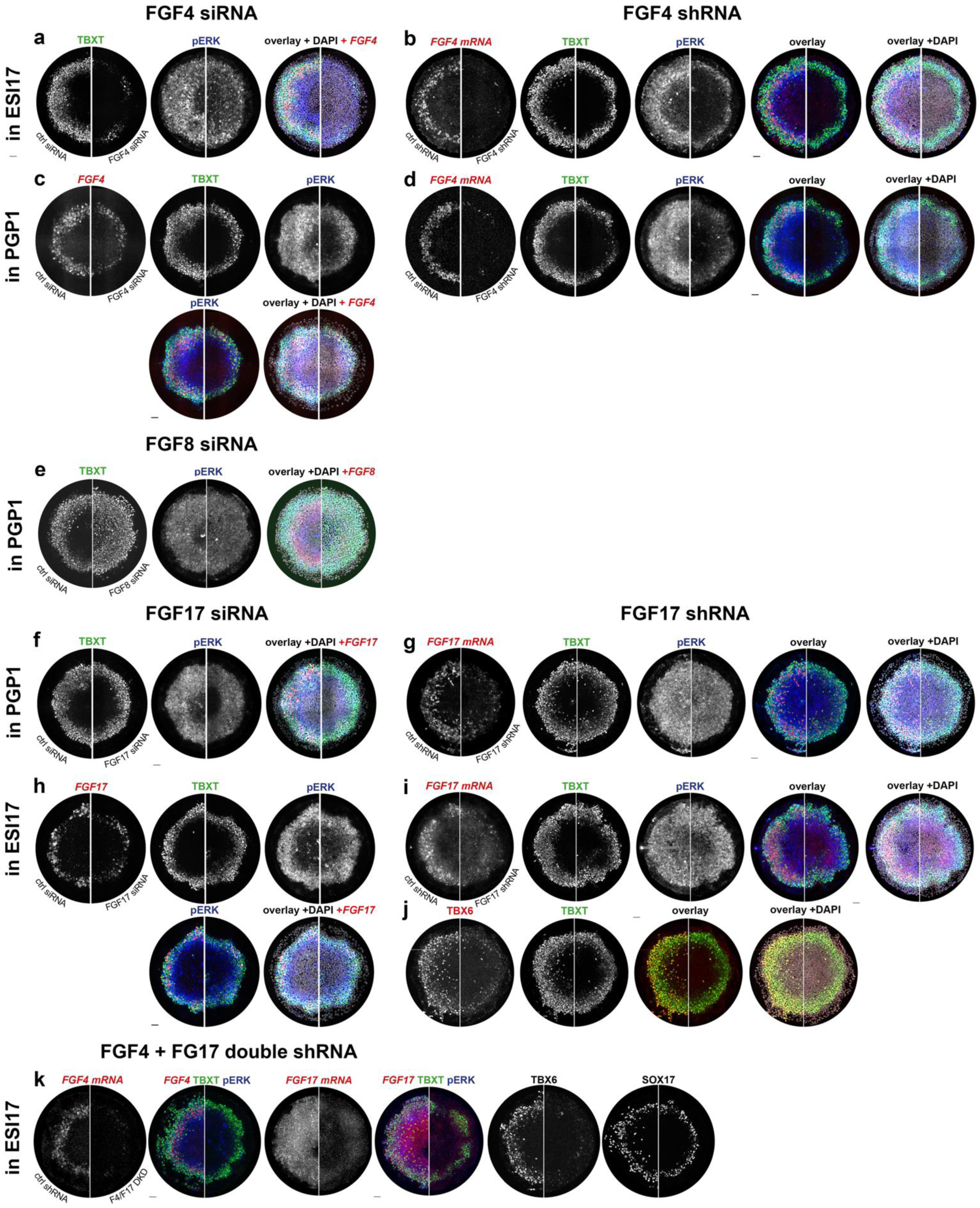
**a)** Additional channels for the colony shown in Fig. 5g. **b)** FGF4 shRNA in ESI17 cells. **c-d)** FGF4 siRNA (c) and shRNA (d) in PGP1 cells. **e-f)** additional channels for Fig. 5h-i. **g**) FGF17 shRNA in ESI17 cells. **h-i)** FGF17 siRNA (h) and shRNA (i) in PGP1 cells. **j)** additional channels for the TBX6 ctrl and FGF17 shRNA in Fig. 5j. **k)** FGF4 + FG17 shRNA double knockdown (stains from multiple colonies).

## References

1. Arnold, S. J. & Robertson, E. J. Making a commitment: cell lineage allocation and axis patterning in the early mouse embryo. Nat. Rev. Mol. Cell Biol. 10, 91–103 (2009).

2. Martyn, I., Kanno, T. Y., Ruzo, A., Siggia, E. D. & Brivanlou, A. H. Self-organization of a human organizer by combined Wnt and Nodal signalling. Nature (2018) doi:10.1038/s41586-018-0150-y.

3. Chhabra, S., Liu, L., Goh, R., Kong, X. & Warmflash, A. Dissecting the dynamics of signaling events in the BMP, WNT, and NODAL cascade during self-organized fate patterning in human gastruloids. PLoS Biol. 17, e3000498 (2019).

4. Etoc, F. et al. A Balance between Secreted Inhibitors and Edge Sensing Controls Gastruloid Self-Organization. Dev. Cell 39, 302–315 (2016).

5. Zhang, Z., Zwick, S., Loew, E., Grimley, J. S. & Ramanathan, S. Embryo geometry drives formation of robust signaling gradients through receptor localization. (2018) doi:10.1101/491290.

6. Heemskerk, I. et al. Rapid changes in morphogen concentration control self-organized patterning in human embryonic stem cells. Elife 8, (2019).

7. Massey, J., et al. WNT ligands stimulate transient signaling in human pluripotent cells and synergize with TGF-β ligands to stimulate sustained signaling during differentiation. bioRxiv 406306 (2018) doi:10.1101/406306.

8. Yang, R. et al. Amnion signals are essential for mesoderm formation in primates. Nat Commun 12, 5126 (2021).

9. Ornitz, D. M. & Itoh, N. The Fibroblast Growth Factor signaling pathway. Wiley Interdiscip Rev Dev Biol 4, 215–266 (2015).

10. LaBonne, C. & Whitman, M. Mesoderm induction by activin requires FGF-mediated intracellular signals. Development 120, 463–472 (1994).

11. Krens, S. F. G., Corredor-Adámez, M., He, S., Snaar-Jagalska, B. E. & Spaink, H. P. ERK1 and ERK2 MAPK are key regulators of distinct gene sets in zebrafish embryogenesis. BMC Genomics 9, 196 (2008).

12. Yao, Y. et al. Extracellular signal-regulated kinase 2 is necessary for mesoderm differentiation. Proc Natl Acad Sci U S A 100, 12759–12764 (2003).

13. Mitrani, E., Gruenbaum, Y., Shohat, H. & Ziv, T. Fibroblast growth factor during mesoderm induction in the early chick embryo. Development 109, 387–393 (1990).

14. Corson, L. B., Yamanaka, Y., Lai, K.-M. V. & Rossant, J. Spatial and temporal patterns of ERK signaling during mouse embryogenesis. Development 130, 4527–4537 (2003).

15. Morgani, S. M. & Hadjantonakis, A.-K. Signaling regulation during gastrulation: Insights from mouse embryos and *in vitro* systems. in Current Topics in Developmental Biology (eds Small, S. & Briscoe, J.) vol. 137 391–431 (Academic Press, 2020).

16. De Simone, A. et al. Control of osteoblast regeneration by a train of Erk activity waves. Nature 590, 129–133 (2021).

17. Marshall, C. J. Specificity of receptor tyrosine kinase signaling: transient versus sustained extracellular signal-regulated kinase activation. Cell 80, 179–185 (1995).

18. Ender, P. et al. Spatiotemporal control of ERK pulse frequency coordinates fate decisions during mammary acinar morphogenesis. Dev Cell 57, 2153–2167.e6 (2022).

19. Aoki, K. et al. Propagating Wave of ERK Activation Orients Collective Cell Migration. Dev Cell 43, 305–317.e5 (2017).

20. Ciruna, B. & Rossant, J. FGF Signaling Regulates Mesoderm Cell Fate Specification and Morphogenetic Movement at the Primitive Streak. Dev. Cell 1, 37–49 (2001).

21. Sun, X., Meyers, E. N., Lewandoski, M. & Martin, G. R. Targeted disruption of Fgf8 causes failure of cell migration in the gastrulating mouse embryo. Genes Dev. 13, 1834–1846 (1999).

22. Fletcher, R. B. & Harland, R. M. The role of FGF signaling in the establishment and maintenance of mesodermal gene expression in Xenopus. Dev Dyn 237, 1243–1254 (2008).

23. Yang, X., Dormann, D., Münsterberg, A. E. & Weijer, C. J. Cell Movement Patterns during Gastrulation in the Chick Are Controlled by Positive and Negative Chemotaxis Mediated by FGF4 and FGF8. Developmental Cell 3, 425–437 (2002).

24. Fürthauer, M., Reifers, F., Brand, M., Thisse, B. & Thisse, C. sprouty4 acts in vivo as a feedback-induced antagonist of FGF signaling in zebrafish. Development 128, 2175–2186 (2001).

25. Tyser, R. C. V. et al. Single-cell transcriptomic characterization of a gastrulating human embryo. Nature 600, 285–289 (2021).

26. Zhai, J. et al. Primate gastrulation and early organogenesis at single-cell resolution. Nature 612, 732–738 (2022).

27. Deng, C. X. et al. Murine FGFR-1 is required for early postimplantation growth and axial organization. Genes Dev 8, 3045–3057 (1994).

28. Yamaguchi, T. P., Harpal, K., Henkemeyer, M. & Rossant, J. fgfr-1 is required for embryonic growth and mesodermal patterning during mouse gastrulation. Genes Dev 8, 3032–3044 (1994).

29. Ciruna, B. G., Schwartz, L., Harpal, K., Yamaguchi, T. P. & Rossant, J. Chimeric analysis of fibroblast growth factor receptor-1 (Fgfr1) function: a role for FGFR1 in morphogenetic movement through the primitive streak. Development 124, 2829–2841 (1997).

30. García-García, M. J. & Anderson, K. V. Essential role of glycosaminoglycans in Fgf signaling during mouse gastrulation. Cell 114, 727–737 (2003).

31. Oki, S., Kitajima, K. & Meno, C. Dissecting the role of Fgf signaling during gastrulation and left-right axis formation in mouse embryos using chemical inhibitors. Dev Dyn 239, 1768–1778 (2010).

32. Feldman, B., Poueymirou, W., Papaioannou, V. E., DeChiara, T. M. & Goldfarb, M. Requirement of FGF-4 for postimplantation mouse development. Science 267, 246–249 (1995).

33. Warmflash, A., Sorre, B., Etoc, F., Siggia, E. D. & Brivanlou, A. H. A method to recapitulate early embryonic spatial patterning in human embryonic stem cells. Nat. Methods 11, 847–854 (2014).

34. Jo, K. et al. Efficient differentiation of human primordial germ cells through geometric control reveals a key role for Nodal signaling. Elife 11, e72811 (2022).

35. Minn, K. T. et al. High-resolution transcriptional and morphogenetic profiling of cells from micropatterned human ESC gastruloid cultures. Elife 9, (2020).

36. Minn, K. T., Dietmann, S., Waye, S. E., Morris, S. A. & Solnica-Krezel, L. Gene expression dynamics underlying cell fate emergence in 2D micropatterned human embryonic stem cell gastruloids. Stem Cell Reports 16, 1210–1227 (2021).

37. Underhill, E. J. & Toettcher, J. E. Control of gastruloid patterning and morphogenesis by the Erk and Akt signaling pathways. Development 150, dev201663 (2023).

38. Loh, K. M. et al. Mapping the Pairwise Choices Leading from Pluripotency to Human Bone, Heart, and Other Mesoderm Cell Types. Cell 166, 451–467 (2016).

39. Moon, K. R. et al. Visualizing structure and transitions in high-dimensional biological data. Nat. Biotechnol. 37, 1482–1492 (2019).

40. Neben, C. L., Lo, M., Jura, N. & Klein, O. D. Feedback regulation of RTK signaling in development. Dev Biol 447, 71–89 (2019).

41. Tyser, R. C. V. & Srinivas, S. Recent advances in understanding cell types during human gastrulation. Seminars in Cell & Developmental Biology 131, 35–43 (2022).

42. Chen, B. et al. Extended culture of 2D gastruloids to model human mesoderm development. bioRxiv 2024.03.21.585753 (2024) doi:10.1101/2024.03.21.585753.

43. Khoa, L. T. P. et al. Visualization of the Epiblast and Visceral Endodermal Cells Using Fgf5-P2A-Venus BAC Transgenic Mice and Epiblast Stem Cells. PLOS ONE 11, e0159246 (2016).

44. Pijuan-Sala, B. et al. A single-cell molecular map of mouse gastrulation and early organogenesis. Nature 566, 490–495 (2019).

45. Morgani, S. M. et al. A Sprouty4 reporter to monitor FGF/ERK signaling activity in ESCs and mice. Developmental Biology 441, 104–126 (2018).

46. Xu, D. & Esko, J. D. Demystifying heparan sulfate-protein interactions. Annu Rev Biochem 83, 129–157 (2014).

47. Habuchi, H. & Kimata, K. Functions of Heparan Sulfate Proteoglycans in Morphogenesis. In Experimental Glycoscience 303–308 (Springer, Tokyo, 2008). doi:10.1007/978-4-431-77922-3_72.

48. Yu, S. R. et al. Fgf8 morphogen gradient forms by a source-sink mechanism with freely diffusing molecules. Nature 461, 533–536 (2009).

49. Benington, L., Rajan, G., Locher, C. & Lim, L. Y. Fibroblast Growth Factor 2-A Review of Stabilisation Approaches for Clinical Applications. Pharmaceutics 12, 508 (2020).

50. Teague, S. et al. Time-integrated BMP signaling determines fate in a stem cell model for early human development. Nat Commun 15, 1471 (2024).

51. Ortiz-Salazar, M. A., Camacho-Aguilar, E. & Warmflash, A. Endogenous Nodal switches Wnt interpretation from posteriorization to germ layer differentiation in geometrically constrained human pluripotent cells. bioRxiv 2024.03.13.584912 (2024) doi:10.1101/2024.03.13.584912.

52. Gurdon, J. B. A community effect in animal development. Nature 336, 772–774 (1988).

53. Nemashkalo, A., Ruzo, A., Heemskerk, I. & Warmflash, A. Morphogen and community effects determine cell fates in response to BMP4 signaling in human embryonic stem cells. Development 144, 3042–3053 (2017).

54. Gattiglio, M., Protzek, M. & Schröter, C. Population-level antagonism between FGF and BMP signaling steers mesoderm differentiation in embryonic stem cells. Biology Open 12, bio059941 (2023).

55. Liu, L. et al. Nodal is a short-range morphogen with activity that spreads through a relay mechanism in human gastruloids. Nat Commun 13, 497 (2022).

56. Isaacs, H. V., Pownall, M. E. & Slack, J. M. eFGF is expressed in the dorsal midline of Xenopus laevis. Int J Dev Biol 39, 575–579 (1995).

57. Christen, B. & Slack, J. M. FGF-8 is associated with anteroposterior patterning and limb regeneration in Xenopus. Dev Biol 192, 455–466 (1997).

58. Hardy, K. M., Yatskievych, T. A., Konieczka, J., Bobbs, A. S. & Antin, P. B. FGF signalling through RAS/MAPK and PI3K pathways regulates cell movement and gene expression in the chicken primitive streak without affecting E-cadherin expression. BMC Dev. Biol. 11, 20–17 (2011).

59. Isaacs, H. V., Pownall, M. E. & Slack, J. M. eFGF regulates Xbra expression during Xenopus gastrulation. The EMBO Journal 13, 4469–4481 (1994).

60. Bulger, E. A., Muncie-Vasic, I., Libby, A. R. G., McDevitt, T. C. & Bruneau, B. G. TBXT dose sensitivity and the decoupling of nascent mesoderm specification from EMT progression in 2D human gastruloids. Development 151, dev202516 (2024).

61. Evans, A. L. et al. Genomic Targets of Brachyury (T) in Differentiating Mouse Embryonic Stem Cells. PLOS ONE 7, e33346 (2012).

62. Nie, Y., Walsh, P., Clarke, D. L., Rowley, J. A. & Fellner, T. Scalable passaging of adherent human pluripotent stem cells. PLoS ONE 9, e88012 (2014).

63. Freeburne, E., et al. Spatial Single Cell Analysis of Proteins in 2D Human Gastruloids Using Iterative Immunofluorescence. Curr Protoc 3, e915 (2023).

64. Sommer, C., Straehle, C., Kothe, U. & Hamprecht, F. A. Ilastik: Interactive learning and segmentation toolkit. in 2011 8th IEEE International Symposium on Biomedical Imaging (ISBI 2011) 230–233 (IEEE, 2011). doi:10.1109/ISBI.2011.5872394.

65. Stringer, C., Wang, T., Michaelos, M. & Pachitariu, M. Cellpose: a generalist algorithm for cellular segmentation. Nat. Methods 18, 100–106 (2021).

66. Wolf, F. A., Angerer, P. & Theis, F. J. SCANPY: large-scale single-cell gene expression data analysis. Genome Biol 19, 15 (2018).

67. Gayoso, A. et al. A Python library for probabilistic analysis of single-cell omics data. Nat Biotechnol 40, 163–166 (2022).

68. van Dijk, D. et al. Recovering Gene Interactions from Single-Cell Data Using Data Diffusion. Cell 174, 716–729.e27 (2018).

69. Street, K. et al. Slingshot: cell lineage and pseudotime inference for single-cell transcriptomics. BMC Genomics 19, 477 (2018).

70. Van den Berge, K., et al. Trajectory-based differential expression analysis for single-cell sequencing data. Nat Commun 11, 1201 (2020).

71. Patro, R., Duggal, G., Love, M. I., Irizarry, R. A. & Kingsford, C. Salmon provides fast and bias-aware quantification of transcript expression. Nat Methods 14, 417–419 (2017).

72. Eswarakumar, V. P., Lax, I. & Schlessinger, J. Cellular signaling by fibroblast growth factor receptors. Cytokine Growth Factor Rev. 16, 139–149 (2005).

